# Investigating biotic, chemical, and physical drivers of periphyton and invertebrate biomass in boreal streams supporting juvenile Atlantic salmon

**DOI:** 10.64898/2026.09.16.752060

**Authors:** Mikael P. Ranta, Shawn J. Leroux, Hannah Adams, Kyleisha J. Foote, Niels van Miltenburg, Nick C. Murphy, Ava J. Hart, Craig F. Purchase

## Abstract

1. Atlantic salmon (*Salmo salar*) are an anadromous fish with a broad distribution from eastern North America to western Europe, with ecological, economic, and cultural significance. Despite their expansive range, wild populations across the North Atlantic have experienced decline. Survival to reproductive age is a management concern, with a focus on identifying factors affecting the juvenile freshwater life stage. Specifically, research is needed to identify potential bottlenecks limiting freshwater productivity.
2. We addressed this research gap with a case study on the Exploits River, Newfoundland – a watershed that receives one of the largest anadromous Atlantic salmon returns in North America. We aimed to identify biotic, chemical, and physical predictors of food resources for juvenile salmon (i.e., invertebrates) and food resources for invertebrates (i.e., periphyton). To test our hypotheses, we collected data on a suite of biotic (e.g., invertebrates and periphyton), chemical (e.g., total dissolved N), and physical (e.g., wetted width) variables at 42 sites within and around this watershed.
3. We observed evidence for relationships between invertebrate biomass and biotic and temporal variables. Specifically, invertebrate biomass was higher at sites with high periphyton biomass that were sampled later in the season. In addition, we observed that periphyton biomass was related to biotic, chemical, and physical variables. Here, we present evidence for bottom-up control, with increased periphyton biomass supporting increased invertebrate biomass.
4. We expect knock-on effects where increased invertebrate biomass results in increased juvenile Atlantic salmon biomass. Our large sample size revealed some predictors of invertebrate biomass and periphyton biomass. However, we found no evidence of correlation between invertebrate biomass and chemical or physical variables. The low productivity of our maritime boreal forest system could be an outlier in expected biotic, chemical, and physical relationships in streams.
5. Our findings provide an important step forward in furthering boreal stream ecology, producing novel invertebrate and periphyton knowledge for watershed managers.

## Introduction

The Atlantic salmon (*Salmo salar*) is an iconic anadromous fish spanning across eastern North America and western Europe (Mather, Parrish, Folt, & DeGraaf, 1998; Rosseland & Kroglund, 2011; Kulmala et al., 2012). Despite their expansive range, wild populations across the North Atlantic have experienced significant decline (Chaput, 2012; Dadswell et al., 2021; ICES, 2023). To slow this decline, management has focused on the survival of salmon to adulthood, or more specifically, on factors affecting the juvenile freshwater life stage (Lai et al., 2024). Anadromous juvenile salmon may spend anywhere from one to eight years in the freshwater environment before migrating to the ocean (Jonsson & Jonsson, 2011). The large variability in the time juveniles spend in fresh water may reflect differences in stream productivity (Poff & Huryn, 1998; Horton, Letcher, Bailey, & Kinnison, 2009), latitudinal gradients (Symons, 1979; Power, 1981), or localized climatic conditions (Jonsson, Jonsson, & Hansen 2005). Research is therefore needed to identify potential bottlenecks limiting freshwater productivity of this species at juvenile life stages.

Salmon production is directly related to invertebrate biomass and production in streams, as invertebrates are the primary food source for juvenile salmon (Keeley & Grant, 1997; Johansen, Elliott, & Klemetsen 2005; Dunlop et al., 2021). Invertebrate biomass and production are largely impacted by a suite of biotic, chemical, and physical stream attributes (Huryn & Wallace, 2000). For example, periphyton (i.e., microbial consortium consisting of algae, bacteria, and organic and inorganic detritus; Wetzel, 1983) in the benthos of streams represents a food source that many invertebrates depend upon for growth and survival (Hart, 1987; Lamberti, Feminella, & Resh, 1987). Consequently, invertebrate biomass and production may be constrained by periphyton (bottom-up), and likewise, periphyton biomass and production may be constrained by invertebrates (top-down; Lamberti & Resh, 1983).

Dissolved nutrients, including nitrogen, within systems may also indirectly influence invertebrate biomass and production through their effects on basal food resources (Cross, Wallace, Rosemond, & Eggert, 2006). These effects may vary among systems, however, with some streams demonstrating positive (Bourassa & Cattaneo, 1998; Robinson & Gessner, 2000), negative (Wang, Robertson, & Garrison, 2007), or no (Simon, Chadwick, Huryn, & Valett, 2010) relationships between stream nutrients and invertebrate biomass and production. Similarly, physical attributes such as stream velocity and water temperature have been shown to have variable effects on invertebrate biomass and production. In some systems, invertebrate assemblages thrive under high flows (Nislow, Folt, & Seandel, 1998; Greenwood & McIntosh, 2010; Caldwell, Rossi, Henery, & Chandra, 2018) and warmer water temperatures (Nelson et al., 2017; Scrine, Jochum, Ólafsson, & O’Gorman, 2017; Jackson et al., 2024). However, opposite relationships have been observed in other systems (Quinn & Hickey, 1990; Cobb, Galloway, & Flannagan, 1992; Hogg & Williams, 1996; Brooks, Haeusler, Reinfelds, & Williams, 2005). These inconsistent relationships suggest that further research is needed to resolve key drivers of invertebrate biomass and production, especially near the northern extent of invertebrate species’ ranges, where drivers remain understudied. Studies in the boreal biome, found between 50°N and 70°N (Hirsch, Trumbore, & Goulden, 2002) and primarily composed of forests dominated by cold-tolerant conifer species (Crosby et al., 2019), may help fill this gap.

Invertebrate genera found in boreal streams are often limited in diversity (Larson & Colbo, 1983); however, these invertebrates are equally, if not more dependent on autochthonous (internally derived) production than those in streams in more productive lower latitudes (Landström, 2015). Forested stream sections in the boreal biome are predominantly comprised of coniferous taxa with low quality litter (Esseen, Ehnström, Ericson, & Sjöberg, 1997; Jonsson, Malmqvist, & Hoffsten, 2001), and as a result, invertebrates may therefore exhibit greater reliance on in situ food resources in these systems. For example, in streams with low in situ primary production, such as those in the boreal biome, invertebrate growth and biomass may be limited with knock-on effects on fish communities that feed on invertebrates. Consequently, juvenile salmon production may be indirectly linked to primary production in streams, as periphyton supports many of the invertebrates that salmon depend upon for growth and survival (Johnston, Perrin, Slaney, & Ward, 1990; Figure 1).

**Figure 1.**
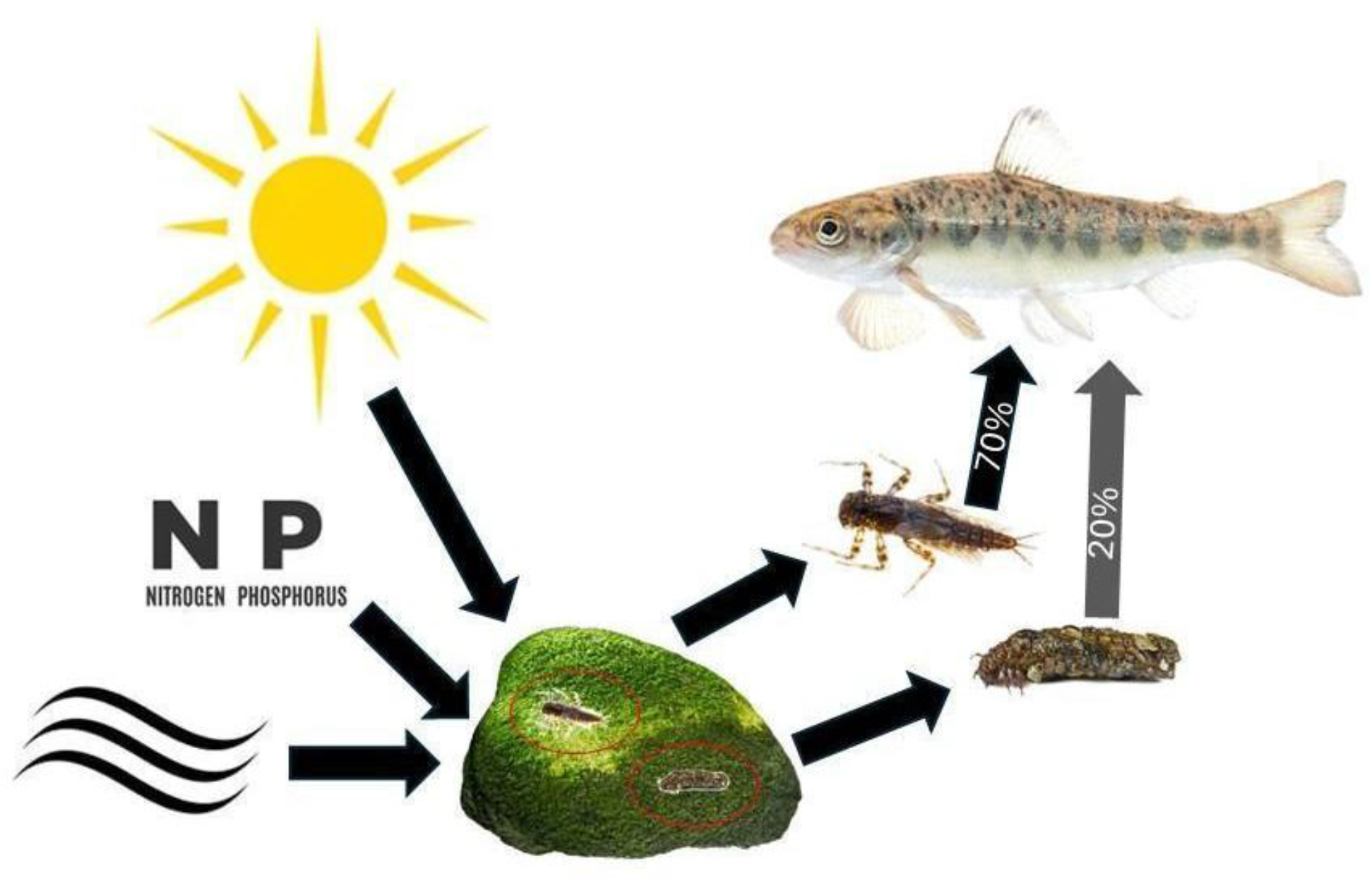
Food web diagram illustrating key bottom-up processes we investigate in an Atlantic salmon river system. Specifically, abiotic factors such as sunlight (promoting photosynthesis), nutrients, and stream velocity may influence periphyton biomass (represented by green growth on the rock). Periphyton biomass supports benthic invertebrates that are in turn preyed upon by juvenile salmon. Differing percentage contribution of invertebrate types toward salmon diet represents potential salmon selectivity or invertebrate limitation. Salmon illustration © Paul van Hoof; Shutterstock (Image ID 1031199370; 1297935142; 1828909271; 2005739255; 2153024879; 2438794217).

Periphyton encompasses a large proportion of primary production occurring in fluvial environments (Li et al., 2022). Periphyton growth and biomass may be influenced by biotic, chemical, and physical factors, which vary across stream and watershed scales. Grazing invertebrates (hereafter, grazers) can be a key biotic constraint on periphyton biomass because many rely on periphyton as a primary food source, and as a result, this reliance may induce significant regulatory effects on periphyton (Kjeldsen, 1996). Regulatory effects of grazers on periphyton may be alleviated, however, when nutrient concentrations are elevated (Liess & Kahlert, 2007). Nutrients, such as nitrogen and phosphorous, are often limiting in streams, and increases in dissolved nutrients consequently have been found to increase periphyton biomass (Lohman, Jones, & Perkins, 1992; Chételat, Pick, Morin, & Hamilton, 1999; Dodds & Smith, 2016). Nutrient increases are not always the main drivers of primary production, however, as primary production in some systems may be more regulated by physical constraints, such as light availability (Bourassa & Cattaneo, 2000; Von Schiller, Martí, Riera, & Sabater, 2007; Hill & Fanta, 2008). Overhanging canopy cover dictates the amount of light available for photosynthesis, and consequently, some streams may experience greater production in the absence of such cover (Lesutiene, Gorokhova, Stankevičienė, Bergman, & Greenberg, 2014), or following leaf abscission in autumn (Hill & Dimick, 2002). Yet, other streams may see more complex interactions, with combinations of light, nutrient availability, or grazer interactions limiting growth; however, neither factor alone serves as the main limiting factor for primary production (Kim & Richardson, 1999; Rosemond, Mulholland, & Brawley 2000; Bernhardt & Likens, 2004; Mallory & Richardson, 2005). Additionally, differences in climate, particularly in the length of the growing season, may influence periphyton growth potential (Day & Henneberg, 2023). Given the variation in regulatory drivers of primary production and standing stocks in streams, there is a need for further research, especially in relatively unproductive systems such as in boreal streams–where less is known about existing drivers.

The identification and characterisation of drivers influencing both invertebrate and periphyton biomass in boreal streams is therefore needed, with results expected to help inform juvenile Atlantic salmon ecology and conservation by identifying bottlenecks associated with juvenile salmon production. To address this research gap, we sampled biotic (invertebrates and periphyton), chemical (e.g., total dissolved N), and physical (e.g., wetted width) components of 42 sites within and around the Exploits River watershed, Newfoundland, Canada in order to characterize invertebrate and periphyton biomass. We tested the hypotheses that a) invertebrate biomass is most influenced by periphyton biomass, in addition to chemical and physical stream attributes, and that b) periphyton biomass is most influenced by invertebrate biomass, in addition to chemical and physical stream attributes. More specifically, we expected sites with higher invertebrate biomass to have i) higher periphyton biomass, ii) higher pH, dissolved nutrients (nitrogen and phosphorous), alkalinity, and conductivity, iii) greater wetted width and stream depth but lower stream velocity, iv) higher Julian date at sampling (even over a narrow sampling window), and v) warmer seasonal water temperatures (i.e., sites with warmer average water temperatures during each defined season). We expected sites with higher periphyton biomass to have vi) higher invertebrate biomass, vii) higher dissolved nutrients (nitrogen and phosphorous), alkalinity, and conductivity, but lower pH, viii) greater wetted width and stream depth but lower stream velocity, ix) higher Julian date at sampling, and x) warmer seasonal water temperatures (see Table 1 for detailed summary and references for predictions).

**Table 1.** Predictor variables used in candidate models for invertebrate and periphyton biomass, including variable names and units, data descriptions, mean values among sites ± standard deviation values across all sites (*n* = 42), predicted relationships with invertebrate and periphyton biomass, and supporting references.

| Variable and units | Description | Mean | Standard deviation | Invertebrate biomass predictions | Periphyton biomass predictions |
| --- | --- | --- | --- | --- | --- |
| Periphyton biomass ($\mu\text{g}/\text{cm}^2$) | Mean biomass of three periphyton samples within sites using chlorophyll *a* as a proxy | 0.05 | 0.08 | Increased food availability via periphyton biomass will lead to increased invertebrate biomass<sup>1,2</sup> | Not used in model |
| Invertebrate biomass ($\text{g}/\text{m}^2$) | Benthic invertebrate biomass was estimated by converting order-level invertebrate densities (individuals/$\text{m}^2$) to biomass using an allometric power-law relationship<sup>3</sup>, with taxon-specific coefficients<sup>4</sup>. | 3.49 | 3.30 | Not used in model | In-stream conditions that support increased invertebrate biomass will also support periphyton growth, allowing for periphyton growth while offsetting losses due to grazing<sup>5,6</sup> |
| Total dissolved nitrogen (TDN; $\text{mg}/\text{L}$) | Mean TDN from three water samples within sites, measured using chemiluminescence detection | 0.20 | 0.06 | Increased total dissolved nitrogen will lead to increased food availability from basal production, which will increase invertebrate biomass<sup>7,8,9</sup> | Increased total dissolved nitrogen will help stimulate growth and increase periphyton biomass<sup>10,11,12,13</sup> |
| pH (pH units) | Mean pH of three water samples (Hanna combo probe) within sites | 6.96 | 0.36 | Certain invertebrate taxa may be more sensitive to lower pH conditions, and as a result, increased pH will support a wider range of taxa and increase invertebrate biomass<sup>14,15</sup> | More acidic conditions will see less invertebrate grazing pressure and increased periphyton biomass; increased pH will have the opposite effect<sup>16,17</sup> |
| Alkalinity ($\text{mg}/\text{L}$ $\text{CaCO}_3$) | Mean alkalinity of three water samples (Lamotte test kit) within sites | 21.84 | 4.45 | Increased alkalinity will expedite detrital decomposition and increase dissolved organic matter available to invertebrates, increasing invertebrate biomass<sup>18</sup> | Increased alkalinity will help stimulate periphyton growth and improve biomass due to an increase in $\text{HCO}_3$ macronutrients<sup>19</sup> |
| Conductivity (EC; $\mu\text{S}/\text{cm}$) | Mean EC of three water samples (Hanna combo probe) within sites | 23.37 | 7.50 | Increased conductivity may reflect an increase in nutrients, which will indirectly increase invertebrate biomass<sup>20</sup> | Increased conductivity may be associated with an increase in nutrients, such as $\text{Ca}^{2+}$, which will improve periphyton biomass<sup>11,21, 22</sup> |
| Wetted width (m) | Mean width of the stream at three intermediate transects within sites | 16.18 | 7.22 | Increased wetted width expands available habitat and potential basal food resources, which will improve invertebrate biomass<sup>23</sup> | Reduced canopy cover from increased wetted width will increase light absorption and improve photosynthesis, increasing periphyton biomass<sup>5,24,25,26</sup> |
| Depth (m) | Mean depth of the stream taken from a range of identical intervals along the three transects within sites | 0.31 | 0.10 | More basal food resources will be found in deeper streams and rivers, in addition to more stable habitat, which will improve invertebrate biomass<sup>27,28</sup> | Increased stream depth will promote periphyton growth and biomass by providing a buffer against physical disturbances and sloughing<sup>7</sup> |
| Velocity (m/s) | Mean velocity rate measured at the three transects within sites | 0.18 | 0.07 | Stable velocity conditions may allow for higher grazing pressure on basal food resources, increasing invertebrate biomass; whereas higher velocity will displace invertebrates from their preferred habitats, decreasing overall invertebrate biomass<sup>7,29</sup> | Higher velocity conditions will limit periphyton growth and may lead to sloughing, reducing periphyton biomass<sup>26,29,30,31,32</sup> |
| Julian date (Day) | Count data representing the accumulated number of days a date represents relative to the beginning of the year | 171.62 | 10.15 | Increased Julian date may be associated with a longer growing season from increased daylight during the time frame sampling occurred (June – July), thereby increasing invertebrate biomass<sup>33,34,35</sup> | Increased Julian date may be associated with a longer growing season from increased daylight and photosynthesis during the time frame sampling occurred (June – July), thereby increasing periphyton biomass<sup>36,37</sup> |
| Water temperature (°C) | Mean water temperatures during spring, summer, fall, and winter seasons recorded using data loggers | 8.81 | 7.15 | Warmer than average water temperatures within spring, summer, fall, or winter seasons will increase biomass of more tolerant invertebrate orders, offsetting losses of more sensitive taxa, leading to overall increased invertebrate biomass<sup>38,39,40</sup> | Warmer than average water temperatures within spring, summer, fall, or winter seasons will lead to more favorable periphyton growth, and result in increased periphyton biomass<sup>41</sup> |

1. Fuller et al., (1986); 2. Kiffney, Richardson, & Bull, (2004); 3. Burgherr & Meyer, (1997); 4. Benke et al., (1999); 5. Kiffney et al., (2003); 6. Wootton, (2012); 7. Bourassa & Cattaneo, (1998); 8. Robinson & Gessner, (2000); 9. Cross et al., (2006); 10. Lohman et al., (1992); 11. Chételat et al., (1999); 12. Lewis Jr. & McCutchan Jr., (2010); 13. Dodds & Smith, (2016); 14. Rosemond, Reice, Elwood, & Mulholland, (1992); 15. Berezina, (2001); 16. Hendry, (1976); 17. Mulholland, Elwood, Palumbo, & Stevenson, (1986); 18. Krueger & Waters, (1983); 19. Dickman, (1973); 20. Reese & Batzer, (2007); 21. Biggs & Price, (1987); 22. Kingsley, Pick, & Hamilton, (2006); 23. Greenwood & McIntosh, (2010); 24. Rounick & Gregory, (1981); 25. Death & Zimmerman, (2005); 26. Warnaars et al., (2007); 27. Kobayashi, Amano, & Nakanishi, (2013); 28. Martínez et al., (2016); 29. Biggs & Stokseth, (1996); 30. Shortreed & Stockner, (1983); 31. Biggs, (1988); 32. Ahn et al., (2013); 33. Clifford, (1972); 34. Shearer et al., (2002); 35. Pasqualini et al., (2023); 36. Biggs, Smith, & Duncan, (1999); 37. Rosemond et al., (2000); 38. Nelson et al., (2017); 39. Scrine et al., (2017); 40. Jackson et al., (2024); 41. Marcarelli & Wurtsbaugh, (2006).

## 2. Methods

### 2.1 Study area and site selection

We established our field study within and around the Exploits River watershed, located in insular Newfoundland, Canada (Figure 2). The drainage basin of the watershed covers 11,272 km^2^ and empties into the Atlantic Ocean at the Bay of Exploits (Scruton et al., 2003). The watershed is composed of many lentic and lotic waterbodies in addition to the main stem (Bourgeois, Murray, & Mercier, 1999). The Exploits River lies in the boreal forest ecozone, within the Central Newfoundland Forest Ecoregion (Department of Fisheries, Forestry, and Agriculture, Forestry & Wildlife Branch, 2021). Heavily forested, the area is dominated by coniferous species such as black spruce (*Picea mariana*) and balsam fir (*Abies balsamea*), as well as sparse deciduous species such as white birch (*Betula papyrifera*). Boglands and barrens are also typical of the area (Cunjak & Newbury, 2005). The geography, climate, flora, and fauna of the ecoregion have been described in more detail by Meades (1990).

**Figure 2.**
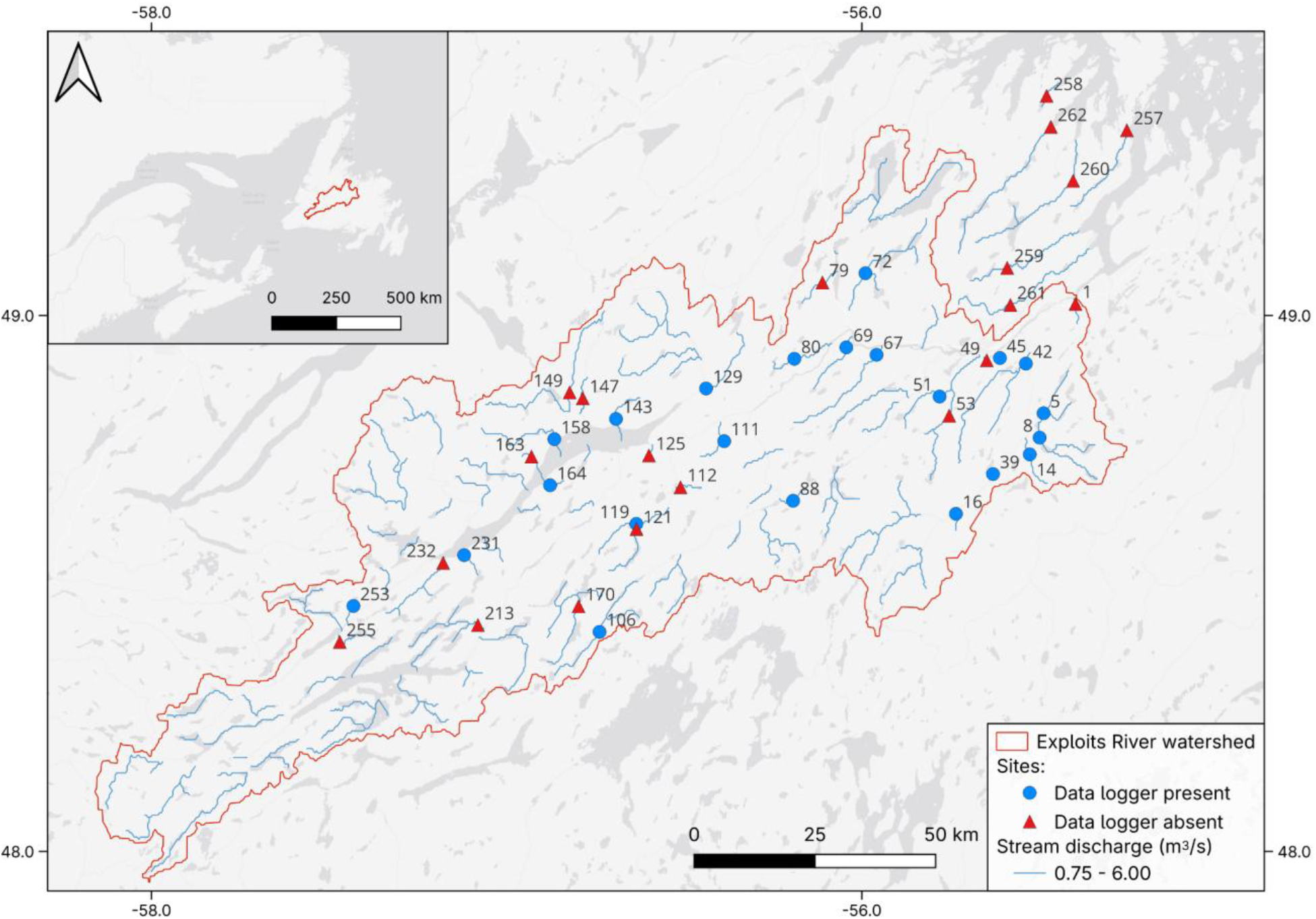
Locations of sampled sites within (red polygon) and around the Exploits River watershed, labelled by site ID in Table S1. Sites with a data logger that measured water temperature are represented by blue circles and sites without a data logger by red triangles. Stream segments within the target discharge range (0.75 - 6.00 m^3^/s) are represented by blue lines. Inset shows the location of the Exploits River watershed (red polygon) in Atlantic Canada. Data sources: Basemap - Esri, HERE, Garmin, © OpenStreetMap contributors, and the GIS User Community; streamlines with discharge – RiverATLAS (Linke et al., 2019).

We used a combination of open access HydroSHEDS data (Lehner, Verdin, & Jarvis, 2008), geological data from the Geoscience Atlas (Newfoundland and Labrador Geological Survey, 2024), and a five m resolution digital elevation model (DEM; Government of Canada, 2024) paired with mean annual discharge filtering (between 0.75 and 6 m^3^/s) to identify 275 potential sites within and around the Exploits River watershed. We narrowed the number of sites for sampling to 42 after filtering by accessibility (< 2 km from road/trail) and the potential for adult anadromous Atlantic salmon to access sites (no impassable barriers). All sites were on different streams, at least 50 m downstream of ponds or waterfalls, at least 50 m upstream of road crossings, developed land, and deforested and thinned forest sections, and more than one km from the ocean in accordance with the Canadian Aquatic Biomonitoring Network (CABIN) protocol (CABIN Field Manual, 2009).

#### 2.1.1 Sampling

We sampled 42 sites within and around the Exploits River watershed, over as short a time frame as was possible, from June 3 – July 6, 2024 (Figure 2; Table S1). We collected data on biotic, chemical and physical features, and recorded the Julian date of sampling (Table S2). To reduce potential bias driven by temporal variation in invertebrate and periphyton responses, we sampled small clusters of sites (∼ 2-4) in different areas of the watershed over time (see Table S1 for specific sampling dates). By following this strategy we avoided having all sites in one area sampled at one time (i.e., not confounding space and time). We established each site over a stream length equivalent of five times the estimated bankfull width as per CABIN guidelines. We measured the length of sites using an open reel tape measure (Crescent Lufkin), and placed marking flags at 25, 50 and 75^th^ percentile lengths. Within each site, we randomly selected riffles for benthic invertebrate and periphyton sampling, and selected the farthest upstream run for water chemistry sampling (CABIN Field Manual, 2009).

### 2.2 Biotic variables

#### 2.2.1 Invertebrate biomass

We mapped each site and divided it into blocks (Figure S1). Each block that contained a riffle was numbered and we used a random number generator (RNG app) to select three blocks for invertebrate collection. We collected benthic invertebrates using a 30.48 x 30.48 cm Surber sampler (hereafter, Surber) with 500 μm mesh. We modified the Surber exteriors with metal “wings” along the base to allow for improved stability by users during invertebrate collection (i.e., users were able to step on wings). To avoid disturbing successive blocks, we started with the furthest downstream block and placed the Surber in a suitable stream bed location (consisting of small cobble or smaller substrate), before gently brushing loose substrate and sediment by hand within Surber confines. After removing all loose substrate, we used a garden trowel (33 x 7 cm) to further prod within the Surber area, which was necessary to collect subsurface invertebrates found in the hyporheic layer (Strommer & Smock, 1989). We then removed collected contents and sifted them through a 500 μm sieve before storing them in 500 mL high-density polyethylene (HDPE) containers with 95% ethanol. We removed large pieces of substrate and other non-invertebrate objects after carefully inspecting them to ensure that invertebrates were not unintentionally discarded.

Given the large number of invertebrates commonly observed in stream sampling, we followed standard protocols to create a representative subsample for invertebrate counting (Hauer & Lamberti, 2007). In the lab, we combined the contents from the three HDPE invertebrate containers collected from each site into a single composite sample (Box S1). We homogenized composite sample contents in a beaker (1,000 mL) using a glass rod (29 x 0.4 cm) and distributed contents among nine labeled sieves ranging from 180-212 µm, visually distributing similar masses to each sieve. We recorded the weights of sieves prior to adding composite sample contents onto them. After we distributed composite sample contents across the sieves, sieves were left to air dry for 20 minutes to allow for excess water and ethanol to evaporate. After drying, we wiped down any noticeable liquid using paper towels, and recorded total weights from each sieve. We determined homogenized content weights (sieve with content weight minus initial sieve weight). We used a random number generator (RNG app) to randomly select three sieves. The remaining contents from the other six sieves were placed in an unsorted 500 mL HDPE container, and the three selected sieves were used for a secondary composite sample. We homogenized the secondary composite sample in a glass beaker (1,000 mL) and filled nine new sieves with secondary composite sample contents following the aforementioned steps. This secondary composite sample was necessary to ensure that at least four total sieves would be fully sorted to reach ≥ 350 invertebrates, when available, so as to meet recommended invertebrate counts associated with satisfactory subsampling accuracy and precision (Glozier, Culp, & Halliwell, 2002). Due to the heterogeneity of contents found within sieves, we sorted at least four sieves. We picked invertebrates from sieves using forceps under a dissecting microscope and preserved invertebrates in vials containing 95% ethanol. We sent the processed invertebrates to BioTech Taxonomy, NB, Canada for order and family-level identification, with invertebrates from 10% of sites (five) additionally verified for quality control by Entomogen Inc., ON, Canada.

For each site, we counted the number of invertebrates from each sorted sieve and calculated the ratio of invertebrates per gram of sample in a sieve (i.e., the invertebrate ratio) by dividing the number of invertebrates collected by the sieve content weight (i.e., individuals/g). We calculated the site-specific median invertebrate ratio from all sorted sieves. We multiplied this value to the total unsorted sample weight (i.e., the summed weight of the six initial sieve contents that were not used for further sorting and all secondary composite sample content weights that were not sorted) to produce an estimate of the number of invertebrates remaining in the unsorted composite sample contents. We added the number of invertebrates sorted to the unsorted estimate to produce an overall estimate of the number of invertebrates in each composite sample. Using this information and the known area of the Surber (0.0929 m^2^), we estimated invertebrate density (individuals/m^2^) for each site.

After receiving taxonomic order and family-level invertebrate information from BioTech, we converted invertebrate density values to biomass density (hereafter, biomass) using published body size data and allometric equations from numerous sources, including the United States Geological Survey (USGS) invertebrate database (Sokol’skaya, 1975; Learner, Lochhead, & Hughes, 1978; Holopainen & Hanski, 1986; Johnston & Cunjak, 1999; Voshell Jr. & Reese, 2002; Vieira et al., 2006; Bouchard Jr., 2009a; Bouchard Jr., 2009b; Edwards, Lauridsen, Armand, Vincent, & Jones, 2009; Goldschmidt & Ramírez Sánchez, 2010; Slobodkin & Bossert, 2010; Coleman, Geisen, & Wall, 2015; Stals, 2015; Vinarski, 2019). From these data, we assigned mean body lengths to each of the orders represented in sorted composite sample contents. We converted the mean length of an individual from each invertebrate order to dry mass using the power law: dry mass = *ax^b^*, where *x* is the length of the organism in mm and *a* and *b* are regression constants (Burgherr & Meyer, 1997). Coefficients for these equations came from Benke, Huryn, Smock, & Wallace (1999), and we used the “all insect” category values for orders where no other coefficients were available (i.e., Acarina, Bivalvia, Cnidaria, Gastropoda, Hirudinea, Neuroptera, Oligochaeta, and Platyhelminthes). From this collective invertebrate data and the Surber area, we derived invertebrate biomass (g/m²) for each site.

#### 2.2.2 Periphyton biomass

At each site we scrubbed periphyton from three total unembedded rocks between 5-10 cm^3^. Individual rocks were collected adjacent to Surber locations (see above and Figure S1), starting with the furthest downstream sampling block. Following Hauer & Lamberti (2007), we used a single coarse bristled toothbrush (22.86 x 5.46 cm) to scrub periphyton from rocks (between 5-10 minutes depending on rock size) and used deionized water to rinse periphyton into a large tray (39.2 x 34.4 cm). We poured periphyton slurries into separate 125 mL containers and placed them in a cooler bag with ice packs. We later (< 5 hours) filtered these slurries through 0.45 μm glass fiber filters (Whatman) using an electric vacuum (VACUUBRAND). Filters were then folded, wrapped in aluminum foil (ALCAN), sealed in plastic freezer bags (Ziploc), and stored in a freezer to minimize chlorophyll degradation until further analysis.

In addition to filtering periphyton, we used aluminum foil to calculate the surface area of each of the rocks we scrubbed, following Hauer & Lamberti (2007). This procedure involved wrapping rocks entirely in aluminum foil at sites, with no overlapping foil. We used the following equation to calculate rock surface area:

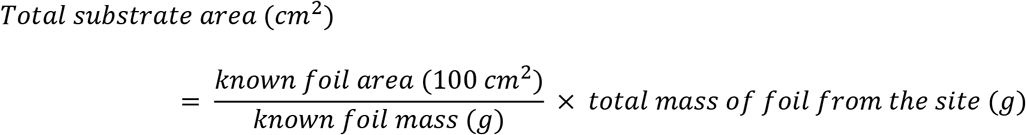

where the mass of aluminum foil required to cover rocks was converted to cm² by referencing the mass of a 100 cm² foil section.

In the lab, we extracted chlorophyll by placing the filters in 90% buffered acetone solution (10% magnesium carbonate) for 12-18 hours while refrigerated at 4 °C and wrapped in aluminum foil (Axler & Owen, 1994; Hauer & Lamberti, 2007). Solutions were then centrifuged at 1850 rpm for 15 minutes at 4 °C (Axler & Owen, 1994). Following centrifuging, we measured chlorophyll *a* content using a SPECTRONIC™ 200 spectrophotometer that measured the absorbance of the sample at 664 nm before and after acidification with 0.1 mL of 0.1 N hydrochloric acid for 90 seconds. We subtracted the absorbance at 750 nm to account for turbidity in the sample and 665 nm to account for the presence of pheophytins as the chlorophyll in the samples degraded (Hauer & Lamberti, 2007). We used the monochromatic equation with an acidification step (see Lorenzen, 1967 and supporting information for details). Following Hauer & Lamberti (2007), we used chlorophyll *a* as a proxy for periphyton biomass. We averaged periphyton biomass values from the three rocks at each site to obtain a single periphyton biomass estimate per site.

### 2.3 Chemical variables

Before collecting invertebrates and periphyton, we used an HDPE cup to collect stream water from the farthest upstream run at each site, which had a minimum surface area of one m^2^. Immediately upon filling the cup with water, we used a pH probe (Hanna HI98129, Hanna Instruments) to measure pH, conductivity, and total dissolved solids. We repeated this to measure three pH, conductivity, and total dissolved solids readings per site and calculated the site mean for each variable. Following readings, we filled three 250 mL amber HDPE bottles with water from the run. We measured alkalinity using water from each bottle with a direct reading titrator (Alkalinity Test Kit 4491-DR-01, Lamotte). At least two measurements were taken at each site, and the average was used to estimate site alkalinity (limited alkalinity tablets led to only two readings taken at most sites).

From the same run, we filled three 250 mL amber HDPE bottles with water, which we later used to measure total dissolved nitrogen (TDN), dissolved organic carbon (DOC), and phosphorous (P) concentrations. Bottles were placed in a cooler bag with ice packs. On the same day, we filtered the water from each bottle through a 0.45 μm polyether sulfone (PES) syringe filter (76479-014, VWR) into pre-combusted and acid-washed 22 mL glass vials for TDN and DOC solutions (three total for both TDN and DOC) and into 15 mL plastic centrifuge tubes (71-1500-B, PROGENE) for P solutions (three total). We preserved each filtered sample to ∼2 pH using three drops of 20% phosphoric acid for TDN and DOC solutions and three drops of 15% nitric acid for P solutions. These solutions were then refrigerated until further lab analysis (methods adapted from Longnecker (2021) and Bowering et al. (2022)). TDN and DOC solutions underwent high temperature combustion (Shimadzu TOC-L), with TDN solutions measured using a chemiluminescence gas analyzer and DOC measured using a non-dispersive infrared gas analyzer. P solutions were measured using optical emission spectroscopy. We removed all resulting data that was below the method detection limit (MDL; TDN = 0.01 mg/L, *n* = 0; DOC = 0.07 mg/L, *n* = 0; P = 0.1 ppm, *n* = 42). All 42 solutions were below the P concentration limit of detection (LOD = 0.1 ppm) and consequently we removed all P data from our analyses.

### 2.4 Physical variables

We measured physical stream characteristics at the 25, 50, and 75th percentile marked lengths of a site, which included wetted width (m), stream depth (m), and stream velocity (m/s). We calculated an average wetted width for each site using the measurements taken at each of the three marked lengths. We used a flow meter (YSI) to measure both stream depth and stream velocity at one m intervals along each transect. Average values were determined for each transect and the three transect values were averaged to determine a single value for each site.

### 2.5 Temporal variable

Our sampling occurred over a 34-day period, and as a result, we investigated whether temporal progression through the summer had an influence on invertebrate and periphyton biomass. To test for this effect, we recorded the Julian date for all sites (i.e., the ordinal number representing the date sampling occurred).

### 2.6 Temperature variables

At 22 sites, we deployed data loggers (HOBO Pendant MX Temp Loggers MX2201; HOBO U20L Water Level Loggers U20L-04) that measure water temperature every hour, and we collected them after almost a full year (Figure 2; Table S1). We secured data loggers on the back of flat rocks ≥ 500 g using zip ties, and placed them in deeper pool sections of streams ≤ 400 m of where sampling occurred.

### 2.7 Statistical analyses

We conducted analyses on two data sets: analyses of biotic, chemical, physical, and temporal predictors from data collected at 42 sites; and analyses of temperature predictors from data collected at 22 sites with temperature loggers. In order to test which measured predictor variables best explained variation in invertebrate and periphyton biomass, we fitted a set of generalized linear models (GLM) with Gamma error distributions and log link functions. We used these models because most of our data were right skewed and did not follow a Gaussian error distribution as revealed by Shapiro-Wilk tests.

#### 2.7.1 Biotic, chemical, physical, and temporal predictors

The suite of models we used were the same for the invertebrate biomass and periphyton biomass responses except the biotic models where periphyton biomass was used as a predictor for invertebrate biomass and invertebrate biomass as a predictor for periphyton biomass (Table S3). Predictor variables used in modelling were as follows: chemical models–total dissolved nitrogen, pH, conductivity, and alkalinity; physical models–wetted width, stream depth, and stream velocity; temporal models–Julian date.

#### 2.7.2 Temperature predictors

Temperature models included average spring, summer, fall, and winter water temperatures as predictors of invertebrate and periphyton biomass (Table S4). Using collected data logger information from 22 sites, we assigned seasons using dates that all loggers were deployed (summer: 6/29/2024-9/21/2024; fall: 9/22/2024-12/20/2024; winter: 12/21/2024-3/19/2025; spring: 3/20/2025-6/18/2025).

#### 2.7.3 Analytical pipeline

Our analytical pipeline consisted of the following steps. First, to avoid overfitting models, we ran a suite of models per prediction category (i.e., biotic, chemical, physical, temporal, and temperature). Second, we had a priori predictions for each individual predictor (see Table 1) and therefore, we fitted models for all combinations of predictors in each category. Third, before fitting models, we used pairwise Pearson’s correlation and variance inflation factor (VIF) analyses to identify collinear (|r| > 0.7) and multi-collinear predictors (VIF > 4). Fourth, we used AIC_c_ (Akaike Information Criterion for small sample size) model selection to identify top ranked models in each category. Following Leroux (2019), we removed models with uninformative parameters. All analyses were conducted in R (ver. 4.5.1; R Core Team, 2025) with packages AICcmodavg (Mazerolle, 2023) and MuMIn (Bartoń, 2024).

Overall, we fitted 25 models with invertebrate biomass as the response and 25 models with periphyton biomass as the response across biotic (1), chemical (15), physical (7), and temporal (1) categories (Table S3). In addition, we fitted five different models with invertebrate biomass as the response and five with periphyton biomass as the response across temperature predictors (Table S4). We identified top models (ΔAIC_c_ = 0), as well as competing models (within ΔAIC_c_ = 2 units away from the top model; Burnham & Anderson 2002) for both response variables.

## 3. Results

### 3.1 Summary

Across our 42 sites, biotic variables of invertebrate biomass and periphyton biomass ranged from 0.33 to 17.44 g/m² and 0.00 to 0.50 μg/cm², respectively. Chemical variables of total dissolved nitrogen and alkalinity ranged between 0.11 and 0.37 mg/L and 14 and 36 mg/L, and physical variables of stream velocity and stream depth ranged between 0.08 and 0.41 m/s and 0.16 and 0.59 m (see Table 1 for mean and standard deviation of all modelled variables).

### 3.2 Invertebrate biomass response

#### 3.2.1 Biotic models

The periphyton biomass model was the top model for explaining variation in invertebrate biomass (*R*^2^ = 0.15, Tables 2 and S5) in the biotic modelling subset. We observed evidence that invertebrate biomass was positively related to periphyton biomass (β = 3.84; SE = 1.68, Figure 3A).

**Figure 3.**
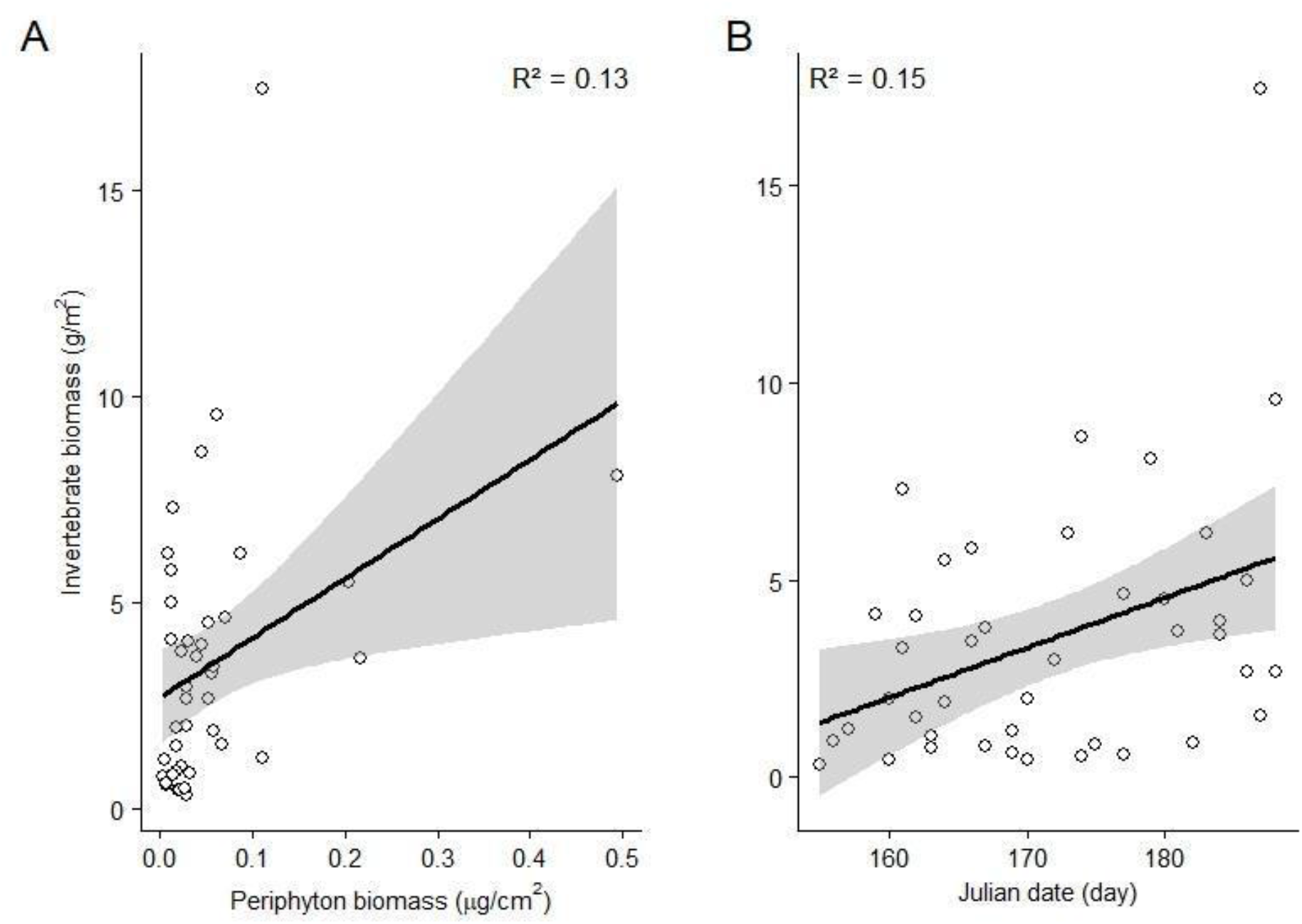
Relationships between (A) invertebrate biomass and periphyton biomass and (B) invertebrate biomass and Julian date. Coefficient of determination (R²); linear regression lines and 95% confidence intervals (shaded) are shown only for statistically significant relationships (*p* < 0.05).

**Table 2.**
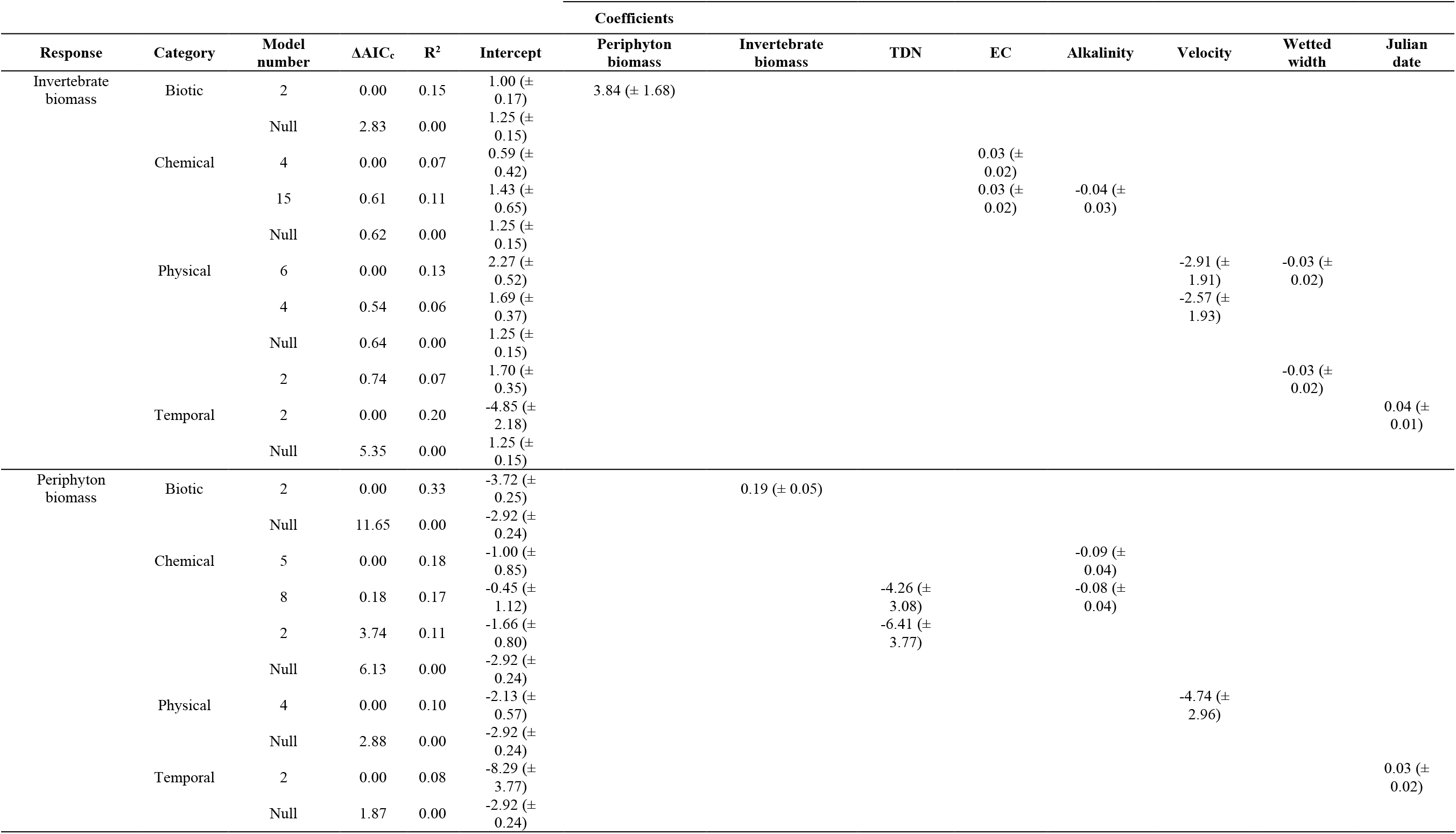
Results of generalized linear models using Gamma error distributions and log link functions to explain invertebrate and periphyton biomass (*n* = 42). We present models within ΔAIC_c_ ≤ 4 of the top model in each category (i.e., biotic, chemical, physical, and temporal). ΔAIC_c_: change in Akaike’s Information Criterion corrected for small sample sizes relative to top model, R^2^: Nagelkerke’s pseudo R^2^ (the proportion of variation in data explained by the model). We have removed models with uninformative variables (see Leroux, 2019 and Table S5 for full AIC table).

#### 3.2.2 Chemical models

We excluded dissolved organic carbon and total dissolved solids from chemical models due to collinearity with selected variables (i.e., total dissolved nitrogen and conductivity). The conductivity model was the top model for explaining variation in invertebrate biomass (*R*^2^ = 0.07, Tables 2 and S5) in the chemical modelling subset; however, it was within 2 ΔAIC_c_ of the null model (ΔAIC_c_ = 0.62), indicating weak to no evidence of a relationship. The conductivity + alkalinity model was a competing model (ΔAIC_c_ = 0.61, *R*^2^ = 0.11), also within 2 ΔAIC_c_ of the null model. We therefore did not find evidence that any of the chemical variables explained the variation in invertebrate biomass. No other competing models ranked above the null model.

#### 3.2.3 Physical models

The stream wetted width + velocity model was the top model for explaining variation in invertebrate biomass (*R*^2^ = 0.13, Tables 2 and S5) in the physical modelling subset; however, it was within 2 ΔAIC_c_ of the null model (ΔAIC_c_ = 0.64), indicating weak to no evidence of a relationship. The velocity (ΔAIC_c_ = 0.54, *R*^2^ = 0.06) and wetted width (ΔAIC_c_ = 0.74, *R*^2^ = 0.07) models were competing models, however, all were within 2 ΔAIC_c_ of the null model. We therefore did not find evidence that any of the physical explanatory variables explained variation in invertebrate biomass. No other competing models ranked above the null model.

#### 3.2.4 Temporal models

Despite purposely restricting the sampling range, the Julian date model was the top model for explaining variation in invertebrate biomass (*R*^2^ = 0.20, Tables 2 and S5) in the temporal modelling subset. We observed evidence that invertebrate biomass was positively related to Julian date (β = 0.04; SE = 0.01, Figure 3B).

#### 3.2.5 Temperature models

The average fall water temperature model was the top model for explaining variation in invertebrate biomass (*R*^2^ = 0.14, Tables 3 and S6) in the temperature modelling subset; however, it was within 2 ΔAIC_c_ of the null model (ΔAIC_c_ = 0.12), indicating weak to no evidence of a relationship. The average spring (ΔAIC_c_ = 1.34, *R*^2^ = 0.09) and winter water temperature (ΔAIC_c_ = 1.68, *R*^2^ = 0.07) models were competing models, however, both were within 2 ΔAIC_c_ of the null model. We therefore did not find evidence that any of the seasonal water temperature explanatory variables explained variation in invertebrate biomass. No other competing models ranked above the null model.

### 3.3 Periphyton biomass response

#### 3.3.1 Biotic models

The invertebrate biomass model was the top biotic model for explaining variation in periphyton biomass (*R*^2^ = 0.33, Tables 2 and S5) in the biotic modelling subset. We observed evidence that periphyton biomass was positively related to invertebrate biomass (β = 0.19; SE = 0.05, Figure 4A).

**Figure 4.**
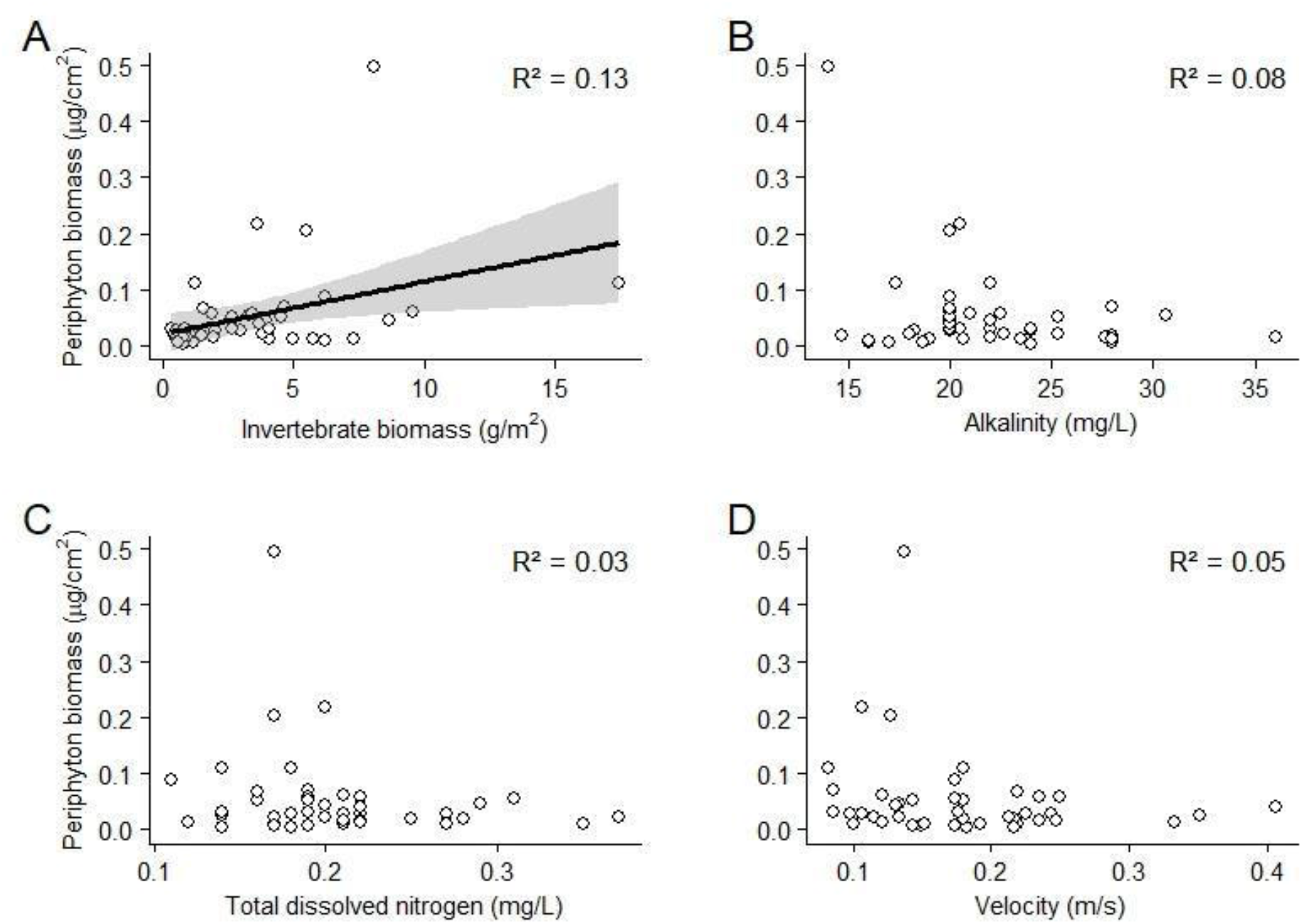
Relationships between (A) periphyton biomass and invertebrate biomass, (B) periphyton biomass and alkalinity, (C) periphyton biomass and total dissolved nitrogen, and (D) periphyton biomass and velocity. Coefficient of determination (R²); linear regression lines and 95% confidence intervals (shaded) are shown only for statistically significant relationships (*p* < 0.05).

#### 3.3.2 Chemical models

The alkalinity model was the top chemical model for explaining variation in periphyton biomass (*R*^2^ = 0.18, Tables 2 and S5) in the chemical modelling subset. The total dissolved nitrogen + alkalinity (ΔAIC_c_ = 0.18, *R*^2^ = 0.17) and ph + alkalinity (ΔAIC_c_ = 1.37, *R*^2^ = 0.18) models were competing models. Periphyton biomass was negatively related to both alkalinity (β = -0.09; SE = 0.04) and total dissolved nitrogen (β = -4.26; SE = 3.08) (Figure 4B and 4C, respectively). No other competing models ranked above the null model.

#### 3.3.3 Physical models

The stream velocity model was the top model for explaining variation in periphyton biomass (*R*^2^ = 0.10, Tables 2 and S5) in the physical modelling subset. We observed evidence that periphyton biomass was negatively related to stream velocity (β = -4.74; SE = 2.96, Figure 4D). No other competing models ranked above the null model.

#### 3.3.4 Temporal models

The Julian date model was the top model for explaining variation in periphyton biomass (*R*^2^ = 0.08, Tables 2 and S5) in the temporal modelling subset; however, it was within 2 ΔAIC_c_ of the null model (ΔAIC_c_ = 1.87), indicating weak to no evidence of a relationship (Tables 2 and S5).

#### 3.3.5 Temperature models

The average summer water temperature model was the top model for explaining variation in periphyton biomass (*R*^2^ = 0.16, Tables 3 and S6) in the temperature modelling subset; however, it was within 2 ΔAIC_c_ of the null model (ΔAIC_c_ = 0.32), indicating weak to no evidence of a relationship. The average fall (ΔAIC_c_ = 0.95, *R*^2^ = 0.07) and winter water temperature (ΔAIC_c_ = 1.89, *R*^2^ = 0.04) models were competing models, however, both were within 2 ΔAIC_c_ of the null model. We therefore did not find evidence that any of the seasonal water temperature explanatory variables explained variation in periphyton biomass. No other competing models ranked above the null model.

**Table 3.** Results of generalized linear models using Gamma error distributions and log link functions to explain invertebrate and periphyton biomass (*n* = 22). We present models within ΔAIC_c_ ≤ 2 of the top model in the temperature category. ΔAIC_c_: change in Akaike’s Information Criterion corrected for small sample sizes relative to top model, R^2^: Nagelkerke’s pseudo R^2^ (the proportion of variation in data explained by the model). See Table S6 for full AIC table.

| Response | Category | Model number | $\Delta AIC_c$ | $R^2$ | Intercept | Coefficients | | | |
| --- | --- | --- | --- | --- | --- | --- | --- | --- | --- |
|  |  |  |  |  |  | Spring water temperature | Summer water temperature | Fall water temperature | Winter water temperature |
| Invertebrate biomass | Temperature | 4 | 0.00 | 0.18 | -2.58 ($\pm 1.92$) | | | 0.58 ($\pm 0.30$) | |
| | | Null | 1.12 | 0.00 | 1.10 ($\pm 0.16$) | | | | |
| Periphyton biomass (mean) | Temperature | 3 | 0.00 | 0.18 | -16.07 ($\pm 7.56$) | | 0.63 ($\pm 0.38$) | | |
| | | 4 | 0.64 | 0.10 | -7.55 ($\pm 3.71$) | | | 0.65 ($\pm 0.59$) | |
| | | Null | 0.98 | 0.00 | -3.39 ($\pm 0.27$) | | | | |

## 4. Discussion

The identification and characterization of drivers influencing both invertebrate and periphyton biomass has been studied in many temperate (Fuller, Roelofs, & Fry, 1986; Marcarelli & Wurtsbaugh, 2006; Greenwood & McIntosh, 2010; Martínez et al., 2016) and inland boreal lotic systems (Veliz, 1999; Raunio & Soininen, 2007; Bechara, Moreau, & Planas, 2011; Burrows, Jonsson, Fältström, Andersson, & Sponseller, 2021), however, a knowledge gap exists for streams in maritime boreal forests, especially within Newfoundland. Central Newfoundland’s unique geology (58% concealed and 24% exposed bedrock in the Exploits River watershed; Government of Newfoundland and Labrador, 2004) may contribute to the low dissolved nutrients found within streams (e.g., phosphorus availability; Bernthal, Armstrong, Nislow, & Metcalfe, 2022) providing a novel environment for researching invertebrate and periphyton biomass drivers. With declining wild Atlantic salmon populations occurring throughout most of Atlantic Canada (DFO, 2024), an emerging fisheries management focus has been placed on identifying constraints on variables known to support the juvenile salmon food base. As such, our goal was to identify potential drivers of invertebrate and periphyton biomass given that both of these biotic compartments directly or indirectly support juvenile Atlantic salmon production. Our results provided evidence for relationships between invertebrate biomass and biotic and temporal variables; and periphyton biomass and biotic, chemical, and physical variables.

We found evidence to support our prediction that invertebrate biomass would increase with periphyton biomass. Allochthonous (externally derived) inputs in the form of deciduous leaf litter are typically less prevalent than coniferous inputs in boreal streams (Landström, 2015), with coniferous litter providing poor energetic value to benthic consumers (Rosset, Barlocher, & Oertli, 1982; Allan & Castillo, 1995). This dynamic was reflected at our sites, with only 2.93% of the invertebrates collected and identified classified within the shredder functional feeding group (i.e., invertebrates that primarily feed on coarse particulate matter; see supporting information for complete invertebrate functional feeding group methods (S2.2.1.a); results in Table S7). Invertebrate growth and biomass in these systems may therefore depend on autochthony, with periphyton serving as a primary energy source. Such invertebrate reliance on periphyton has been demonstrated in both second and third order streams, with reductions in periphyton biomass having direct implications on invertebrate biomass and densities (Fuller et al., 1986; Heaston, Kaylor, & Warren, 2018). A meta-analysis across 89 studies likewise found that when periphyton abundance was reduced, grazer invertebrate density and growth decreased (Feminella & Hawkins, 1995). Our findings of a positive relationship between invertebrate biomass and periphyton biomass are consistent with these results. Across the sites we sampled, mean periphyton biomass was 0.05 μg/cm². This value is relatively low when compared with values found in other boreal waterbodies, which suggests that streams in the Exploits River watershed are not very productive. For example, a mean value of 0.71 μg/cm² was reported in a Finnish river (Raunio & Soininen, 2007) and streams within a Swedish watershed averaged 0.85 μg/cm² (Landström, 2015). Similarly, in New Brunswick streams, periphyton biomass collected from artificial tiles deployed in July and August and collected in August and September ranged from 0.03-2.49 μg/cm² (Erdozain, Kidd, Kreutzweiser, & Sibley, 2018). Although our mean periphyton biomass value falls within this range, the majority of our periphyton biomass values are at the lower end of this distribution. The low periphyton biomass we found across our sites may stem from a variety of factors (e.g., chemical or physical), however, invertebrates at sites with higher periphyton biomass appeared to capitalize on the available food resources, resulting in greater overall invertebrate biomass in those areas.

We found evidence to support our prediction of a positive relationship between invertebrate biomass and Julian date of sampling (over an approximate one month period). The seasonal relationship between invertebrate biomass and Julian date has been well documented, with in-stream invertebrate density and biomass peaks observed in late June to mid-July (Pearson & Franklin, 1968; Clifford, 1972; Cellot, 1996; Barton & Farmer, 1997; Shearer, Hayes, & Stark, 2002, Pasqualini, Majdi, & Brauns, 2023), or early fall (Chadwick & Canton, 1983; Sagar & Glova, 1992). Our sampling timeframe was conducted from early June to early July, and as a result, it is possible that the invertebrate populations we sampled at many sites had not reached peak biomass yet. As the summer progresses, an increasing number of invertebrate eggs hatch, resulting in a variety of instars occupying the benthos (Chadwick & Canton, 1983). This community-wide life cycle progression may therefore explain the positive relationship we found. Alternatively, although Julian date represents an ordinal value that aligns with calendar date, it may serve as a proxy for other environmental conditions, even over a narrow sampling window (e.g., warming water temperatures or lengthened daylight during the summer). For example, in our data set, we observed significant relationships between Julian date and conductivity, stream depth, and stream velocity (see Figures S2 and S3 in supporting information).

Contrary to our other predictions, our data did not provide evidence to support relationships between invertebrate biomass and total dissolved nitrogen, phosphorus, stream velocity, or seasonal water temperatures. We expected to see bottom-up effects between dissolved nutrients and invertebrate biomass, as previously observed in headwater streams in North Carolina, USA (Greenwood & Rosemond, 2005), with sites with higher total dissolved nitrogen and phosphorous concentrations having increased periphyton biomass, and consequently increased invertebrate biomass. Although dissolved nutrients within our sites were low, we still expected to find evidence of a trophic bridge connecting nutrient availability to invertebrate biomass, as nutrient availability can influence invertebrate community dynamics in boreal streams (Bergfur, Johnson, Sandin, Goedkoop, & Nygren, 2007). However, as phosphorus was below detection limits at all our sites, we could not test for a relationship between phosphorus and invertebrate biomass. Additionally, the streams we sampled seem to have very low primary production (see above), thus, it is possible that indirect effects from chemical variables are not detectable in this watershed due to low variability in nitrogen concentrations (e.g., TDN: 0.11-0.37 mg/L). This could also be indicative of a phosphorus-limited system (Elser et al., 2007). Regardless, the relationship between nutrients and invertebrates may become more apparent with the inclusion of sites with higher nutrient concentrations.

Stream velocity and seasonal water temperatures can have differing influences on invertebrate communities (Quinn & Hickey, 1990; Hogg & Williams, 1996; Nislow et al., 1998; Scrine et al., 2017). Our results are consistent with work done on streams in New Zealand, but opposite to studies from Arizona and England. In New Zealand, invertebrate biomass among sites did not differ when stream flows were within 20X median flow rates (Quinn & Hickey, 1990). Our results reflect this dynamic, as summer flows during sampling remained relatively stable. If the sites we sampled had been subject to flash-flooding, as seen in Arizona streams (Grimm & Fisher, 1989), we would have expected a significant, negative relationship between velocity and invertebrate biomass. Similarly, the effects of warming water temperatures on invertebrate assemblages may become more pronounced when greater variation within the temperature gradient exists. In a meta-analysis across 50 English streams, sites increased in temperature by 2.1-2.9 °C in winter and 1.1-1.5 °C in summer over a 26-year period, resulting in noticeable invertebrate assemblage shifts in chalk streams (Durance & Ormerod, 2009). Seasonal water temperature variation within the sites we sampled was relatively low in three out of four seasons (3.75-10.03% CV), and data loggers were deployed for around a year. Given our sampling timeframe, it is possible that water temperature influences on invertebrate biomass were subtle, and therefore may only become detected through longer-term monitoring or with further invertebrate sampling across the other three seasons.

The positive relationship between invertebrate and periphyton biomass may reflect the importance of periphyton to invertebrate diets. If the biotic communities among the sites we sampled would have been top-down controlled, we would have expected a negative relationship between periphyton and invertebrate biomass. Consistent with our prediction of a bottom-up controlled dynamic, we found evidence to support a positive relationship between periphyton and invertebrate biomass. This finding could be considered unexpected as the effects of invertebrate grazing have been found to have immediate implications on periphyton growth and biomass (Hillebrand, 2002), with increased invertebrate biomass often resulting in decreased periphyton biomass (Feminella, Power, & Resh, 1989; Boston & Hill, 1991; Jones, Young, Eaton, & Moss, 2002). From the sites that we sampled, only 6.13% of invertebrates that were taxonomically identified were categorized as scraper invertebrates. This low percentage may offer explanation for the overall positive relationship between periphyton and invertebrate biomass if most of the invertebrates throughout the watershed were not primarily consuming periphyton, but rather using periphyton as habitat.

In some instances, in-stream conditions may also produce a favorable environment where both periphyton and invertebrate biomass may simultaneously increase, and periphytic losses from grazing are offset by rapid growth. For example, in the South Fork Pysht River, Washington, USA, 100-300 m sections of overhanging alders were removed and replaced with conifer seedlings, resulting in 13-fold periphyton biomass and 7-fold invertebrate density increases (Wootton, 2012). Similarly, sections of headwater streams in British Columbia, Canada, were experimentally clear cut, significantly increasing both periphyton biomass and invertebrate density (Kiffney, Richardson, & Bull, 2003). Future work could measure primary production along with standing biomass, as in our existing study, in order to better parse out the relationship between invertebrates and periphyton in boreal streams. Primary production can be measured readily by placing artificial tiles in streams for a period and later collecting them to measure accumulated periphyton biomass (see example in MacSween, Leroux, & Oakes (2019)). Adding electrical current around some tiles can also isolate the impacts of biotic vs chemical/physical variables on primary production in streams (see example in Kurle & Cardinale (2011)).

A further explanation for the positive relationship between periphyton and invertebrate biomass may be found within the cellular makeup of periphyton. As periphyton growth expands, films of diatoms are replaced by macroalgae, and invertebrate families that previously grazed on periphyton are replaced by filterer and collector invertebrates, which use the dense structure as a form of habitat (Dudley, Cooper, & Hemphill 1986; Tonkin, Death, & Barquín 2014). Although overall periphyton biomass was relatively low within the Exploits River watershed, invertebrate reliance on periphyton as both a source of food and habitat may be expected, as 26.48% of invertebrates collected and identified were classified as filterer/collectors. The relatively high abundance of filterer and collector invertebrates across our sites may therefore have contributed towards the positive relationship we observed.

In most nutrient-limited streams, increased nitrogen and phosphorous concentrations are associated with increased periphyton biomass (Lohman et al., 1992; Chételat et al., 1999; Dodds & Smith, 2016). We expected our results to align with this prediction, as total dissolved nitrogen levels averaged 0.20 mg/L across the sites we sampled, and all phosphorous solutions collected were below method detection limits – evidence of a nutrient-depleted environment. The total dissolved nitrogen average from our solutions was additionally much lower than average concentrations found in other boreal watersheds. In a smaller Newfoundland watershed, total dissolved nitrogen levels averaged 0.84 mg/L (Adams & Leroux, 2024), and in Sweden 0.77 mg/L (Långberg, 2024). Contrary to our expectation, we found evidence to support a negative relationship between periphyton biomass and total dissolved nitrogen. One explanation for why this may have occurred is that sites with higher total dissolved nitrogen may have also had other variables that were associated with lower periphyton biomass. Bourassa & Cattaneo (1998) found that in southern Quebec streams periphyton biomass was not related to nutrient increases, with other factors such as grazing invertebrates proposed to have greater controlling influences on periphyton growth. Similarly, other variables we sampled may have had a greater influence on periphyton biomass accrual. For example, we found evidence to support negative relationships between periphyton biomass and alkalinity and periphyton biomass and stream velocity. Alkalinity is typically not found to have a significant impact on periphyton biomass, although an increase in HCO_3_ (bicarbonate) macronutrients associated with alkalinity may stimulate periphytic growth (Dickman, 1973). The negative relationship we found most likely demonstrates that HCO_3_ macronutrients are not limiting periphyton growth within the Exploits River watershed.

Conversely, the negative relationship between periphyton biomass and stream velocity has been well documented (Shortreed & Stockner, 1983; Biggs & Stokseth, 1996; Bourassa & Cattaneo, 1998). Primary reasoning for such a dynamic may be attributed to growth potential. In higher flow environments, periphyton growth is lessened and complete detachment via sloughing may occur (Biggs, 1988; Warnaars, Hondzo, & Power 2007). Such a result was supported in a mesocosm experiment, where periphyton biomass increased with stream velocity until critical values were reached (0.20-0.50 m/s; Ahn et al., 2013). Although none of the sites we sampled exceeded a velocity of 0.41 m/s, our study watershed is very wet, receiving ∼ 1000-1500 mm of precipitation per year (Government of Newfoundland and Labrador, 2020), with much of this falling and accumulating as snow from December to April. Consequently, it is possible that higher flow conditions earlier in the year following the annual spring snowmelt greatly reduced periphyton biomass at many sites. In addition, the geology across most of the sites we sampled consisted of exposed or nearly exposed bedrock (Government of Newfoundland and Labrador, 2004). This exposure would greatly reduce the absorptive buffering capacity along streams and promote higher discharge during high water events. Biggs & Stokseth (1996) found that periphyton biomass was highest at velocities < 0.30 m/s in a New Zealand river and lowest > 0.70 m/s in a Norwegian river. Consequently, we may expect that the streams we sampled that had higher velocities most likely were at velocities above favorable periphyton growth thresholds prior to when sampling occurred. Periphyton biomass accrual at these sites following reduced stream velocities as the summer progressed would likely remain lower than periphyton biomass in more buffered and stable sections of streams, offering a potential explanation for the negative relationship we found.

In many smaller order streams, canopy cover dictates light availability, and consequently may constrain periphyton growth (Stovall, Keeton, & Kraft 2009; Warren, Keeton, Bechtold, & Rosi-Marshall, 2013). If canopy cover cannot be accurately estimated, stream wetted width may serve as a proxy. As stream wetted width increases, canopy cover decreases (Vannote, Minshall, Cummins, Sedell, & Cushing 1980), as trees cannot indefinitely grow laterally over streams. As such, we used wetted width as a way to measure potential light limitation of periphyton within our sites, with a prediction that increased wetted width would result in increased periphyton biomass. Our data, however, failed to support this prediction. All of the sites we sampled had a wetted width of at least five m and many sites had less than one m of overhanging riparian vegetation or tree cover, resulting in predominantly unattenuated sunlight. This lack of shading could explain why light was not limiting photosynthetic growth in periphyton at the sites we sampled. Sections of streams in the headwaters of the sites we sampled may have experienced more pronounced periphyton-light induced relationships, however, targeted sampling in locations less than five m wetted width with prominent overhanging cover would need to be done to test for this potential effect.

In this study, we highlight potential variables constraining invertebrate and periphyton biomass in a nutrient-depleted, boreal watershed. Further work may be warranted, however, to better understand how invertebrate and periphyton biomass may influence tertiary consumer dynamics. Here, we present evidence for bottom-up control, with increased periphyton biomass supporting increased invertebrate biomass. Consequently, we may expect knock-on effects where increased invertebrate biomass results in increased juvenile Atlantic salmon biomass. Sampling to determine juvenile salmon biomass within these streams is therefore needed to confirm the direction of energy exchange and identify the degree of importance aquatic invertebrates contribute towards juvenile salmon diet, length, weight, and ages. Atlantic salmon populations across Atlantic Canada have continued to decline (Dadswell et al., 2021; ICES, 2023). By linking indirect influences from periphyton biomass and direct influences from invertebrate biomass to juvenile salmon biomass, fisheries managers may be able to identify areas within watersheds that would most benefit from habitat modifications or enhancement projects. The estimated increase in juvenile salmon could then potentially lead to an increase in adult returns. Such results were found in Washington, USA, where habitat improvements and flow restoration in Salmon Creek led to measurable increases in invertebrate and juvenile rainbow trout (*Oncorhynchus mykiss*) populations, resulting in improved juvenile recruitment to the adult population and increased adult returns (ISRP, 2025). Future work within the Exploits River watershed would be needed to implement such improvements, with short and long term monitoring to facilitate learning by doing. Our present findings, however, provide an important step forward in furthering boreal stream ecology, producing novel invertebrate and periphyton knowledge for watershed managers.

## Acknowledgements

Funding for this research was provided by a) the Department of Fisheries and Oceans Canada Aquatic Ecosystems Restoration Fund to the Environment Resources Management Association, b) the Foundation for Conservation of Atlantic Salmon to Memorial University of Newfoundland, c) Memorial University of Newfoundland, d) Canada Foundation for Innovation, and e) the Natural Sciences and Engineering Research Council of Canada. We want to extend our appreciation towards T. Paul, K. Sheppard, and D. Ryan for assistance with the project and L. Rowsell, R. Pearcey, C. Hanley, R. Wagle, and E. Wilson for technical assistance in the field and lab.

## Author Contribution Statement

Conceptualization, developing methods: MR, SL, CP, HA, KF, NVM, NM. Conducting the research: MR, SL, CP, KF, NVM, NM. Data analysis: MR, SL, CP, NVM, NM, AH. Data interpretation: MR, SL, CP, NVM, NM, KF, HA, AH. Preparation of figures and tables: MR, SL, CP, KF. Writing: MR, SL, CP, KF, NVM, NM, HA, AH.

## Data Availability Statement

The data that support the findings of this study are available in the following Figshare repository: https://doi.org/10.6084/m9.figshare.33752380

## References

Adams, H. & Leroux, S. J. (2024). Integrating field data and a meta-ecosystem model to study the effects of multiple terrestrial disturbances on small stream ecosystem function. Ecosystems 27(7), 951–968. DOI: 10.1007/s10021-024-00932-x.

Ahn, C. H., Song, H. M., Lee, S., Oh, J. H., Ahn, H., Park, J. R., Lee, J. M., & Joo, J. C. (2013). Effects of water velocity and specific surface area on filamentous periphyton biomass in an artificial stream mesocosm. Water, 5(4), 1723–1740. DOI: 10.3390/w5041723.

Allan, J. D. & Castillo, M. M. (1995) Stream ecology: structure and function of running waters (2nd ed.). Springer Netherlands, Netherlands.

Axler, R. P., & Owen, C. J. (1994). Measuring chlorophyll and phaeophytin: Whom should you believe? Lake and Reservoir Management, 8(2), 143–151. DOI: 10.1080/07438149409354466.

Barbour, M. T., Gerritsen, J., Snyder, B. D., & Stribling, J. B. (1999) Rapid bioassessment protocols for use in streams and wadeable rivers: Periphyton, benthic macroinvertebrates, and fish. EPA 841-OB-99-002. United States Environmental Protection Agency, Washington, DC.

Barton, D. R., & Farmer, M. E. D. (1997). The effects of conservation tillage practices on benthic invertebrate communities in headwater streams in southwestern Ontario, Canada. Environmental Pollution 96(2), 207–215. DOI: 10.1016/0269-7491(97)00020-1.

Bartoń, K. (2024) MuMIn: Multi-model inference. R package version 1.48–4.

Bechara, J. A., Moreau, G., & Planas, D. (1992). Top-down effects of brook trout (*Salvelinus fontinalis*) in a boreal forest stream. Canadian Journal of Fisheries and Aquatic Sciences, 49(10), 2093–2103. DOI: 10.1139/f92-233.

Benke, A. C., Huryn, A. D., Smock, L. A., & Wallace, J. B. (1999). Length-mass relationships for freshwater macroinvertebrates in North America with particular reference to the southeastern United States. Journal of the North American Benthological Society, 18(3), 308–343.

Berezina, N. A. (2001). Influence of ambient pH on freshwater invertebrates under experimental conditions. Russian Journal of Ecology, 32(5), 343–351. DOI: 10.1023/A:1011978311733.

Bergfur, J., Johnson, R. K., Sandin, L., Goedkoop, W., & Nygren, K. (2007). Effects of nutrient enrichment on boreal streams: Invertebrates, fungi and leaf-litter breakdown. Freshwater Biology, 52(8), 1618–1633. DOI: 10.1111/j.1365-2427.2007.01770.x.

Bernhardt, E. S., & Likens, G. E. (2004). Controls on periphyton biomass in heterotrophic streams. Freshwater Biology, 49(1), 14–27. DOI: 10.1046/j.1365-2426.2003.01161.x.

Bernthal, F. R., Armstrong, J. D., Nislow, K. H., & Metcalfe, N. B. (2022). Nutrient limitation in Atlantic salmon rivers and streams: Causes, consequences, and management strategies. Aquatic Conservation: Marine and Freshwater Ecosystems, 32(6), 1073–1091. DOI: 10.1002/aqc.3811.

Biggs, B. J. F. (1988). Artificial substrate exposure times for periphyton biomass estimates in rivers. New Zealand Journal of Marine and Freshwater Research, 22(4), 507–515. DOI: 10.1080/00288330.1988.9516321.

Biggs, B. J. F., & Price, G. M. (1987). A survey of filamentous algal proliferations in New Zealand rivers. New Zealand Journal of Marine and Freshwater Research, 21(2), 175–191. DOI: 10.1080/00288330.1987.9516214.

Biggs, B. J. F., & Stokseth, S. (1996). Hydraulic habitat suitability for periphyton in rivers. River Research and Applications, 12(2–3), 251–261. DOI: 10.1002/(SICI)1099-1646(199603)12:2/3<251::AID-RRR393>3.0.CO;2-X.

Biggs, B. J. F., Smith, R. A., & Duncan, M. J. (1999). Velocity and sediment disturbance of periphyton in headwater streams: Biomass and metabolism. Journal of the North American Benthological Society, 18(2), 222–241. DOI: 10.2307/1468462.

Boston, H. L., & Hill, W. R. (1991). Photosynthesis–light relations of stream periphyton communities. Limnology and Oceanography, 36(4), 644–656. DOI: 10.4319/lo.1991.36.4.0644.

Bouchard, R. W. (2009a) Neuroptera (Spongillaflies). Chapter 9. In: Guide to aquatic invertebrate families of Mongolia: Identification manual for students, citizen monitors, and aquatic resource professionals. University of Minnesota, St Paul, MN.

Bouchard, R. W. (2009b) Lepidoptera. Chapter 11. In: Guide to aquatic invertebrate families of Mongolia: Identification manual for students, citizen monitors, and aquatic resource professionals. University of Minnesota, St Paul, MN.

Bourassa, N., & Cattaneo, A. (1998). Control of periphyton biomass in Laurentian streams (Quebec). Journal of the North American Benthological Society, 17(4), 420–429. DOI: 10.2307/1468363.

Bourassa, N., & Cattaneo, A. (2000). Responses of a lake outlet community to light and nutrient manipulation: Effects on periphyton and invertebrate biomass and composition. Freshwater Biology, 44(4), 629–639. DOI: 10.1046/j.1365-2427.2000.00610.x.

Bourgeois, C. E., Murray, J. & Mercier, V. (1999) Status of the Exploits River stock of Atlantic salmon (*Salmo salar* L.) in 1998. Canadian Stock Assessment Secretariat Research Document 99/82, 27 pp.

Bowering, K. L., Edwards, K. A., Wiersma, Y. F., Billings, S. A., Warren, J., Skinner, A., & Ziegler, S. E. (2023). Dissolved organic carbon mobilization across a climate transect of mesic boreal forests is explained by air temperature and snowpack duration. Ecosystems, 26(1), 55–71. DOI: 10.1007/s10021-022-00741-0.

Brooks, A. J., Haeusler, T. I. M., Reinfelds, I., & Williams, S. (2005). Hydraulic microhabitats and the distribution of macroinvertebrate assemblages in riffles. Freshwater Biology, 50(2), 331–344. DOI: 10.1111/j.1365-2427.2004.01322.x.

Burgherr, P., & Meyer, E. I. (1997). Regression analysis of linear body dimensions vs. dry mass in stream macroinvertebrates. Archiv für Hydrobiologie, 101–112. DOI: 10.1127/archiv-hydrobiol/139/1997/101.

Burnham, K. P., & Anderson, D. R. (2002). *Model selection and multimodel inference: A practical information-theoretic approach* (2nd ed.). New York, NY: Springer. DOI: 10.1007/b97636.

Burrows, R. M., Jonsson, M., Fältström, E., Andersson, J., & Sponseller, R. A. (2021). Interactive effects of light and nutrients on stream algal growth modified by forest management in boreal landscapes. Forest Ecology and Management, 492, 119212. DOI: 10.1016/j.foreco.2021.119212.

Caldwell, T. J., Rossi, G. J., Henery, R. E., & Chandra, S. (2018). Decreased streamflow impacts fish movement and energetics through reductions to invertebrate drift body size and abundance. River Research and Applications, 34(8), 965–976. DOI: 10.1002/rra.3340.

Chadwick, J. W., & Canton, S. P. (1983). Comparison of multiplate and Surber samplers in a Colorado mountain stream. Journal of Freshwater Ecology, 2(3), 287–292.

Chaput, G. (2012). Overview of the status of Atlantic salmon (*Salmo salar*) in the North Atlantic and trends in marine mortality. ICES Journal of Marine Science, 69(9), 1538–1548. DOI: 10.1093/icesjms/fss013.

Chételat, J., Pick, F. R., Morin, A., & Hamilton, P. B. (1999). Periphyton biomass and community composition in rivers of different nutrient status. Canadian Journal of Fisheries and Aquatic Sciences, 56(4), 560–569. DOI: 10.1139/f98-197.

Clifford, H. F. (1972). A years’ study of the drifting organisms in a brown-water stream of Alberta, Canada. Canadian Journal of Zoology 50(7), 975–983. DOI: 10.1139/z72-130.

Cobb, D. G., Galloway, T. D., & Flannagan, J. F. (1992). Effects of discharge and substrate stability on density and species composition of stream insects. Canadian Journal of Fisheries and Aquatic Sciences, 49(9), 1788–1795. DOI: 10.1139/f92-198.

Coleman, D. C., Geisen, S. & Wall, D. H. (2024) Soil fauna: occurrence, biodiversity, and roles in ecosystem function. In: Soil microbiology, ecology and biochemistry (eds E. Paul & S. Frey), pp. 131–159. Elsevier. DOI: 10.1016/B978-0-12-822941-5.00005-3.

Collins, J. J. (2017) Platyhelminthes. Current Biology, 27(7), R252–R256. DOI: 10.1016/j.cub.2017.02.016.

Crosby, A. D., Bayne, E. M., Cumming, S. G., Schmiegelow, F. K. A., Dénes, F. V., & Tremblay, J. A. (2019). Differential habitat selection in boreal songbirds influences estimates of population size and distribution. Diversity and Distributions, 25(12), 1941–1953. DOI: 10.1111/ddi.12991.

Cunjak, R. A. & Newbury, R. W. (2005) Atlantic coast rivers of Canada. In: Rivers of North America (eds AC Benke & CE Cushing), pp. 939–980. Elsevier Academic Press, San Diego, CA.

Cross, W. F., Wallace, J. B., Rosemond, A. D., & Eggert, S. L. (2006). Whole-system nutrient enrichment increases secondary production in a detritus-based ecosystem. Ecology, 87(6), 1556–1565. DOI: 10.1890/0012-9658(2006)87[1556:WNEISP]2.0.CO;2.

Dadswell, M., Spares, A., Reader, J., McLean, M., McDermott, T., Samways, K., & Lilly, J. (2021). The decline and impending collapse of the Atlantic Salmon (*Salmo salar*) population in the North Atlantic Ocean: A review of possible causes. Reviews in Fisheries Science & Aquaculture, 30(2), 215–258. DOI: 10.1080/23308249.2021.1937044.

Day, N. K., & Henneberg, M. F. (2023) Investigation of potential factors controlling benthic algae in the upper White River Basin, Colorado, 2018–21. US Geological Survey Scientific Investigations Report 2023–5009, 30 pp. DOI: 10.3133/sir20235009.

Death, R. G., & Zimmermann, E. M. (2005). Interaction between disturbance and primary productivity in determining stream invertebrate diversity. Oikos, 111(2), 392–402. DOI: 10.1111/j.0030-1299.2005.13799.x.

Department of Fisheries, Forestry, and Agriculture. (2021). Zone 3 forest management plan 2022–2026. St. John’s, Canada: Government of Newfoundland and Labrador.

Department of Fisheries and Oceans Canada. (2024). Canadian Science Advisory Secretariat Science Advisory Report (Report No. 2025/007). Ottawa, Canada: Fisheries and Oceans Canada.

Dhooria, M. S. (2016) Fundamentals of applied acarology. Springer, Singapore, 470 pp.

Dickman, M. (1973). Changes in periphytic algae following bicarbonate additions to a small stream. Journal of the Fisheries Research Board of Canada, 30(12), 1882–1884. DOI: 10.1139/f73-318.

Dodds, W. K., & Smith, V. H. (2016). Nitrogen, phosphorus, and eutrophication in streams. Inland Waters, 6(2), 155–164. DOI: 10.5268/IW-6.2.909.

Dudley, T. L., Cooper, S. D., & Hemphill, N. (1986). Effects of macroalgae on a stream invertebrate community. Journal of the North American Benthological Society, 5(2), 93–106. DOI: 10.2307/1467864.

Dunlop, K., Eloranta, A. P., Schoen, E., Wipfli, M., Jensen, J. L., Muladal, R., & Christensen, G. N. (2021). Evidence of energy and nutrient transfer from invasive pink salmon (*Oncorhynchus gorbuscha*) spawners to juvenile Atlantic salmon (*Salmo salar*) and brown trout (*Salmo trutta*) in northern Norway. Ecology of Freshwater Fish, 30(2), 270–283. DOI: 10.1111/eff.12582.

Durance, I., & Ormerod, S. J. (2009). Trends in water quality and discharge confound long-term warming effects on river macroinvertebrates. Freshwater Biology, 54(2), 388–405. DOI: 10.1111/j.1365-2427.2008.02112.x.

Edwards, F. K., Lauridsen, R. B., Armand, L., Vincent, H. M., & Jones, J. I. (2009). The relationship between length, mass and preservation time for three species of freshwater leeches (Hirudinea). Fundamental and Applied Limnology, 173(4), 321–327. DOI: 10.1127/1863-9135/2009/0173-0321.

Elser, J. J., Bracken, M. E. S., Cleland, E. E., Gruner, D. S., Harpole, W. S., Hillebrand, … Smith, J. E. (2007). Global analysis of nitrogen and phosphorus limitation of primary producers in freshwater, marine and terrestrial ecosystems. Ecology Letters, 10(12), 1135–1142. DOI: 10.1111/j.1461-0248.2007.01113.x.

Erdozain, M., Kidd, K., Kreutzweiser, D., & Sibley, P. (2018). Linking stream ecosystem integrity to catchment and reach conditions in an intensively managed forest landscape. Ecosphere, 9(5), e02278. DOI: 10.1002/ecs2.2278.

Esseen, P. A., Ehnström, B., Ericson, L., & Sjöberg, K. (1997). Boreal forests. Ecological Bulletins, 46, 16–47. DOI: 10.2307/20113207.

Feminella, J. W., & Hawkins, C. P. (1995). Interactions between stream herbivores and periphyton: A quantitative analysis of past experiments. Journal of the North American Benthological Society, 14(4), 465–509. DOI: 10.2307/1467536.

Feminella, J. W., Power, M. E., & Resh, V. H. (1989). Periphyton responses to invertebrate grazing and riparian canopy in three northern California coastal streams. Freshwater Biology, 22(3), 445–457. DOI: 10.1111/j.1365-2427.1989.tb01117.x.

Fuller, R. L., Roelofs, J. L., & Fry, T. J. (1986). The importance of algae to stream invertebrates. Journal of the North American Benthological Society, 5(4), 290–296. DOI: 10.2307/1467481.

Glozier, N., Culp, J., & Halliwell, D. (2002). Revised guidance for sample sorting and sub-sampling protocols for EEM benthic invertebrate community surveys. Saskatoon, Canada: National Water Research Institute, Environment Canada.

Goldschmidt, T., & Ramirez Sanchez, M. M. (2020). Introduction and keys to Neotropical water mites. Spixiana, 43(2), 203–303.

Government of Canada. (2024). High Resolution Digital Elevation Model (HRDEM) – CanElevation Series. Retrieved May 11, 2024, from https://open.canada.ca/data/en/dataset/957782bf-847c-4644-a757-e383c0057995.

Government of Newfoundland and Labrador. (2004). Water quality station profile: Sutherlands Pond outflow at Forest Access Road (NF02YN0044). Retrieved August 24, 2026, from https://www.canal.gov.nl.ca/root/main/station_details_e.asp?envirodat=NF02YN0044.

Government of Newfoundland and Labrador. (2020). Hydrology and Climate of Newfoundland. Municipal and Community Affairs. Retrieved July 28, 2026, from https://www.gov.nl.ca/mca/nl/.

Greenwood, J. L., & Rosemond, A. D. (2005). Periphyton response to long-term nutrient enrichment in a shaded headwater stream. Canadian Journal of Fisheries and Aquatic Sciences, 62(9), 2033–2045. DOI: 10.1139/f05-117.

Greenwood, M. J., & McIntosh, A. R. (2010). Low river flow alters the biomass and population structure of a riparian predatory invertebrate. Freshwater Biology, 55(10), 2062–2076. DOI: 10.1111/j.1365-2427.2010.02462.x.

Grimm, N. B., & Fisher, S. G. (1989). Stability of periphyton and macroinvertebrates to disturbance by flash floods in a desert stream. Journal of the North American Benthological Society, 8(4), 293–307. DOI: 10.2307/1467493.

Hart, D. D. (1987). Experimental studies of exploitative competition in a grazing stream insect. Oecologia, 73(1), 41–47. DOI: 10.1007/BF00376975.

Hauer, F. R. & Lamberti, G. A. (2007) Methods in stream ecology (2nd ed.). Academic Press, Burlington, MA.

Heaston, E. D., Kaylor, M. J., & Warren, D. R. (2018). Aquatic food web response to patchy shading along forested headwater streams. Canadian Journal of Fisheries and Aquatic Sciences, 75(12), 2211–2220. DOI: 10.1139/cjfas-2017-0464.

Hendry, G. R. (1976). Effects of pH on the growth of periphytic algae in artificial stream channels (Report No. IR 25/76). Oslo, Norway: SNSF Project.

Hill, W. R., & Dimick, S. M. (2002). Effects of riparian leaf dynamics on periphyton photosynthesis and light utilisation efficiency. Freshwater Biology, 47(7), 1245–1256. DOI: 10.1046/j.1365-2427.2002.00837.x.

Hill, W. R., & Fanta, S. E. (2008). Phosphorus and light colimit periphyton growth at subsaturating irradiances. Freshwater Biology, 53(2), 215–225. DOI: 10.1111/j.1365-2427.2007.01885.x.

Hillebrand, H. (2002). Top-down versus bottom-up control of autotrophic biomass—a meta-analysis on experiments with periphyton. Journal of the North American Benthological Society, 21(3), 349–369. DOI: 10.2307/1468275.

Hirsch, A. I., Trumbore, S. E., & Goulden, M. L. (2002). Direct measurement of the deep soil respiration accompanying seasonal thawing of a boreal forest soil. Journal of Geophysical Research: Atmospheres, 107(D23), WFX-2. DOI: 10.1029/2001JD000921.

Hogg, I. D., & Williams, D. D. (1996). Response of stream invertebrates to a global-warming thermal regime: An ecosystem-level manipulation. Ecology, 77(2), 395–407. DOI: 10.2307/2265617.

Holopainen, I. J., & Hanski, I. (1986). Life history variation in Pisidium (Bivalvia: Pisidiidae). Ecography, 9(2), 85–98. DOI: 10.1111/j.1600-0587.1986.tb01195.x.

Horton, G. E., Letcher, B. H., Bailey, M. M., & Kinnison, M. T. (2009). Atlantic salmon (*Salmo salar*) smolt production: The relative importance of survival and body growth. Canadian Journal of Fisheries and Aquatic Sciences, 66(3), 471–483. DOI: 10.1139/F09-005.

Huryn, A. D., & Wallace, J. B. (2000). Life history and production of stream insects. Annual Review of Entomology, 45(1), 83–110. DOI: 10.1146/annurev.ento.45.1.83.

ICES. (2023). *Working group on North Atlantic Salmon (WGNAS)* (ICES Scientific Reports 5(41)). Copenhagen, Denmark: International Council for the Exploration of the Sea.

Independent Scientific Review Panel (ISRP). (2025). Habitat retrospective report: Review and synthesis of progress and challenges in Columbia River Basin Fish and Wildlife Program habitat protection and restoration projects (Report No. ISRP 2025-2). Portland, OR: Northwest Power and Conservation Council.

Jackson, M. C., Friberg, N., Moliner Cachazo, L., Clark, D. R., Mutinova, P. T., O’Gorman, E. J., … Woodward, G. (2024). Regional impacts of warming on biodiversity and biomass in high latitude stream ecosystems across the Northern Hemisphere. Communications Biology, 7(1), 316. DOI: 10.1038/s42003-024-05936-w.

Johansen, M., Elliott, J. M., & Klemetsen, A. (2005). Relationships between juvenile salmon, *Salmo salar* L., and invertebrate densities in the River Tana, Norway. Ecology of Freshwater Fish, 14(4), 331–343. DOI: 10.1111/j.1600-0633.2005.00107.x.

Johnston, N. T., Perrin, C. J., Slaney, P. A., & Ward, B. R. (1990). Increased juvenile salmonid growth by whole-river fertilization. Canadian Journal of Fisheries and Aquatic Sciences, 47(5), 862–872. DOI: 10.1139/f90-099.

Johnston, T. A., & Cunjak, R. A. (1999). Dry mass–length relationships for benthic insects: A review with new data from Catamaran Brook, New Brunswick, Canada. Freshwater Biology, 41(4), 653–674. DOI: 10.1046/j.1365-2427.1999.00400.x.

Jones, J. I., Young, J. O., Eaton, J. W., & Moss, B. (2002). The influence of nutrient loading, dissolved inorganic carbon and higher trophic levels on the interaction between submerged plants and periphyton. Journal of Ecology, 90(1), 12–24. DOI: 10.1046/j.0022-0477.2001.00620.x.

Jonsson, B. & Jonsson, N. (2011) Smolts and smolting. In: Ecology of Atlantic salmon and brown trout: Habitat as a template for life histories (eds B. Jonsson & N. Jonsson), pp. 211–245. Springer, Dordrecht. DOI: 10.1007/978-94-007-1189-1_5.

Jonsson, N., Jonsson, B. & Hansen, L.P. (2005) Does climate during embryonic development influences parr growth and age of seaward migration in Atlantic salmon (*Salmo salar*) smolts? Canadian Journal of Fisheries and Aquatic Sciences, 62(11), 2502–2508. DOI: 10.1139/f05-154.

Jonsson, M., Malmqvist, B., & Hoffsten, P. O. (2001). Leaf litter breakdown rates in boreal streams: Does shredder species richness matter? Freshwater Biology, 46(2), 161–171. DOI: 10.1046/j.1365-2427.2001.00655.x.

Keeley, E. R., & Grant, J. (1997). Allometry of diet selectivity in juvenile Atlantic salmon (*Salmo salar*). Canadian Journal of Fisheries and Aquatic Sciences, 54(8), 1894–1902. DOI: 10.1139/f97-096.

Kiffney, P. M., Richardson, J. S., & Bull, J. P. (2003). Responses of periphyton and insects to experimental manipulation of riparian buffer width along forest streams. Journal of Applied Ecology, 40(6), 1060–1076. DOI: 10.1046/j.1365-2664.2003.00750.x.

Kiffney, P. M., Richardson, J. S., & Bull, J. P. (2004). Establishing light as a causal mechanism structuring stream communities in response to experimental manipulation of riparian buffer width. Journal of the North American Benthological Society, 23(3), 542–555. DOI: 10.1899/0887-3593(2004)023<0542:ELAACM>2.0.CO;2.

Kim, M. A., & Richardson, J. S. (2000). Effects of light and nutrients on grazer–periphyton interactions. In L. M. Darling (Ed.), Proceedings of a Conference on the Biology and Management of Species and Habitats at Risk, Kamloops, B.C., 15–19 February 1999 (Vol. 2, pp. 497–502). B.C. Ministry of Environment, Lands and Parks & University College of the Cariboo.

Kingsley, M., Pick, F. R., & Hamilton, P. B. (2006). Periphyton biomass and composition of British Columbia salmonid rivers in relation to nutrient and discharge levels. Internationale Vereinigung für theoretische und angewandte Limnologie: Verhandlungen, 29(3), 1389–1398. DOI: 10.1080/03680770.2005.11902910.

Kjeldsen, K. (1996). Regulation of algal biomass in a small lowland stream: Field experiments on the role of invertebrate grazing, phosphorus and irradiance. Freshwater Biology, 36(3), 535–546. DOI: 10.1046/j.1365-2427.1996.00111.x.

Kobayashi, S., Amano, K., & Nakanishi, S. (2013). Riffle topography and water flow support high invertebrate biomass in a gravel-bed river. Freshwater Science, 32(3), 706–718. DOI: 10.1899/12-080.1.

Krueger, C. C., & Waters, T. F. (1983). Annual production of macroinvertebrates in three streams of different water quality. Ecology, 64(4), 840–850. DOI: 10.2307/1937207.

Kulmala, S., Haapasaari, P., Karjalainen, T. P., Kuikka, S., Pakarinen, T., Parkkila, K., Romakkaniemi, A. & Vuorinen, P. J. (2013) Ecosystem services provided by the Baltic salmon: A regional perspective to the socio-economic benefits associated with a keystone species. In: Socio-economic importance of ecosystem services in the Nordic countries: Synthesis in the context of The Economics of Ecosystems and Biodiversity (TEEB), pp. 266–276. Nordic Council of Ministers, Copenhagen.

Kurle, C. M., & Cardinale, B. J. (2011). Ecological factors associated with the strength of trophic cascades in streams. Oikos, 120(12), 1897–1908. DOI: 10.1111/j.1600-0706.2011.19465.x.

Lai, S. Y., Pálsson, A., Guðbergsson G., Jónsson, I. R., Ólafsson, J. S., & Bárðarson, H. (2024). The prey availability and diet of juvenile Atlantic salmon (*Salmo salar* L.) in low-productivity rivers in northern Europe. Journal of Fish Biology, 105(1), 72–84. DOI: 10.1111/jfb.15757.

Lamberti, G. A., & Resh, V. H. (1983). Stream periphyton and insect herbivores: An experimental study of grazing by a caddisfly population. Ecology, 64(5), 1124–1135. DOI: 10.2307/1937823.

Lamberti, G. A., Feminella, J. W., & Resh, V. H. (1987). Herbivory and intraspecific competition in a stream caddisfly population. Oecologia, 73(1), 75–81. DOI: 10.1007/BF00376980.

Landström, E. (2015) Resource use by macroinvertebrates within boreal stream food webs. Master’s Thesis, Umeå University.

Långberg, A. (2024) Stream periphyton biomass along a rural-urban gradient: An analysis of factors influencing periphyton biomass in streams within and around Umeå. Bachelor’s Thesis, Umeå University.

Larson, D. J. & Colbo, M. H. (1983) The aquatic insects: Biogeographic considerations. In: Biogeography and ecology of the island of Newfoundland (ed. G. R. South), pp. 593–677. Dr W. Junk Publishers, The Hague.

Learner, M. A., Lochhead, G., & Hughes, B. D. (1978). A review of the biology of British Naididae (Oligochaeta) with emphasis on the lotic environment. Freshwater Biology, 8(4), 357–375. DOI: 10.1111/j.1365-2427.1978.tb01457.x.

Lehner, B., Verdin, K., & Jarvis, A. (2008). New global hydrography derived from spaceborne elevation data. *Eos*, Transactions American Geophysical Union, 89(10), 93–94. DOI: 10.1029/2008eo100001.

Leroux, S. J. (2019). On the prevalence of uninformative parameters in statistical models applying model selection in applied ecology. PloS One, 14(2), e0206711. DOI: 10.1371/journal.pone.0206711.

Lesutienė, J., Gorokhova, E., Stankevičienė, D., Bergman, E., & Greenberg, L. (2014). Light increases energy transfer efficiency in a boreal stream. PLoS One, 9(11), e113675. DOI: 10.1371/journal.pone.0113675.

Lewis, W. M., Jr., & McCutchan, J. H., Jr. (2010). Ecological responses to nutrients in streams and rivers of the Colorado mountains and foothills. Freshwater Biology, 55(9), 1973–1983. DOI: 10.1111/j.1365-2427.2010.02431.x.

Li, N., Hao, Y., Sun, H., Wu, Q., Tian, Y., Mo, J., … Guo, J. (2022). Distribution and photosynthetic potential of epilithic periphyton along an altitudinal gradient in Jue River (Qinling Mountain, China). Freshwater Biology, 67(10), 1761–1773. DOI: 10.1111/fwb.13973.

Liess, A., & Kahlert, M. (2007). Gastropod grazers and nutrients, but not light, interact in determining periphytic algal diversity. Oecologia, 152(1), 101–111. DOI: 10.1007/s00442-006-0636-4.

Linke, S., Lehner, B., Ouellet Dallaire, C., Ariwi, J., Grill, G., Anand, M., Beames, P., Burchard-Levine, V., Maxwell, S., Moidu, H., Tan, F., & Thieme, M. (2019). *Global* hydro-environmental sub-basin and river reach characteristics at high spatial resolution. Scientific Data, 6(1), 283. DOI: 10.1038/s41597-019-0300-6.

Lohman, K., Jones, J. R., & Perkins, B. D. (1992). Effects of nutrient enrichment and flood frequency on periphyton biomass in northern Ozark streams. Canadian Journal of Fisheries and Aquatic Sciences, 49(6), 1198–1205. DOI: 10.1139/f92-135.

Longnecker, K. (2022). Collecting samples for Dissolved Organic Carbon (DOC) analysis (Version 1). protocols.io. Woods Hole, MA: Woods Hole Oceanographic Institution. 10.17504/protocols.io.b6xrrfm6.

Lorenzen, C. J. (1967). Determination of chlorophyll and pheo-pigments: Spectrophotometric equations 1. Limnology and Oceanography, 12(2), 343–346. DOI: 10.4319/lo.1967.12.2.0343.

MacSween, J., Leroux, S. J., & Oakes, K. D. (2019). Cross-ecosystem effects of a large terrestrial herbivore on stream ecosystem functioning. Oikos, 128(1), 135–145. DOI: 10.1111/oik.05331.

Mallory, M. A., & Richardson, J. S. (2005). Complex interactions of light, nutrients and consumer density in a stream periphyton–grazer (tailed frog tadpoles) system. Journal of Animal Ecology, 74(6), 1020–1028. DOI: 10.1111/j.1365-2656.2005.01000.x.

Marcarelli, A. M., & Wurtsbaugh, W. A. (2006). Temperature and nutrient supply interact to control nitrogen fixation in oligotrophic streams: An experimental examination. Limnology and Oceanography, 51(5), 2278–2289. DOI: 10.4319/lo.2006.51.5.2278.

Martínez, A., Basaguren, A., Larrañaga, A., Molinero, J., Pérez, J., Sagarduy, M., & Pozo, J. (2016). Differences in water depth determine leaf-litter decomposition in streams: implications on impact assessment reliability. Knowledge and Management of Aquatic Ecosystems, (417), 23. DOI: 10.1051/kmae/2016010.

Mather, M. E., Parrish, D. L., Folt, C. L., & DeGraaf, R. M. (1998). Integrating across scales: Effectively applying science for the successful conservation of Atlantic salmon (*Salmo salar*). Canadian Journal of Fisheries and Aquatic Sciences, 55(S1), 1–8. DOI: 10.1139/d98-000.

Mazerolle, M. J. (2023) AICcmodavg: Model selection and multimodel inference based on (Q)AIC(c). R package version 2.3–4.

Meades, S. J. (1990) Natural regions of Newfoundland and Labrador. Protected Areas Association, St John’s, Newfoundland and Labrador.

Ministry of Environment. (2009). The Canadian Aquatic Biomonitoring Network: Field Manual (Version 1.0). Victoria, Canada: Resources Information Standards Committee.

Mulholland, P. J., Elwood, J. W., Palumbo, A. V., & Stevenson, R. J. (1986). Effect of stream acidification on periphyton composition, chlorophyll, and productivity. Canadian Journal of Fisheries and Aquatic Sciences, 43(10), 1846–1858. DOI: 10.1139/f86-229.

Nelson, D., Benstead, J. P., Huryn, A. D., Cross, W. F., Hood, J. M., Johnson, P. W., … Ólafsson, J. S. (2017). Shifts in community size structure drive temperature invariance of secondary production in a stream-warming experiment. Ecology, 98(7), 1797–1806. DOI: 10.1002/ecy.1857.

Newfoundland and Labrador Geological Survey. (2024). Newfoundland and Labrador GeoScience Atlas Online. Retrieved from https://geoatlas.gov.nl.ca.

Nislow, K. H., Folt, C., & Seandel, M. (1998). Food and foraging behavior in relation to microhabitat use and survival of age-0 Atlantic salmon. Canadian Journal of Fisheries and Aquatic Sciences, 55(1), 116–127. DOI: 10.1139/f97-222.

Oksanen, L., Fretwell, S. D., Arruda, J., & Niemela, P. (1981). Exploitation ecosystems in gradients of primary productivity. The American Naturalist, 118(2), 240–261. DOI: 10.1086/283817.

Oliveira, A. L. H., & Nessimian, J. L. (2010) Spatial distribution and functional feeding groups of aquatic insect communities in Serra da Bocaina streams, southeastern Brazil. Acta Limnologica Brasiliensia, 22(4), 424–441. DOI: 10.4322/actalb.2011.007.

Pasqualini, J., Majdi, N., & Brauns, M. (2023). Effects of incomplete sampling on macroinvertebrate secondary production estimates in a forested headwater stream. Hydrobiologia, 850(14), 3113–3124. DOI: 10.1007/s10750-023-05238-y.

Pearson, W. D., & Franklin, D. R. (1968). Some factors affecting drift rates of Baetis and Simuliidae in a large river. Ecology 49(1), 75–81. DOI: 10.2307/1933562.

Poff, N. L., & Huryn, A. D. (1998). Multi-scale determinants of secondary production in Atlantic salmon (*Salmo salar*) streams. Canadian Journal of Fisheries and Aquatic Sciences, 55(S1), 201–217. DOI: 10.1139/d98-013.

Power, G. (1981) Stock characteristics and catches of Atlantic salmon (*Salmo salar*) in Quebec, and Newfoundland and Labrador in relation to environmental variables. Canadian Journal of Fisheries and Aquatic Sciences, 38(12), 1601–1611. DOI: 10.1139/f81-210.

Quinn, J. M., & Hickey, C. W. (1990). Characterization and classification of benthic invertebrate communities in 88 New Zealand rivers in relation to environmental factors. New Zealand Journal of Marine and Freshwater Research, 24(3), 387–409. DOI: 10.1080/00288330.1990.9516432.

Raunio, J., & Soininen, J. (2007). A practical and sensitive approach to large river periphyton monitoring: comparative performance of methods and taxonomic levels. Boreal Environment Research, 12(1), 55–63.

Reese, E. G., & Batzer, D. P. (2007). Do invertebrate communities in floodplains change predictably along a river’s length? Freshwater Biology, 52(2), 226–239. DOI: 10.1111/j.1365-2427.2006.01678.x.

Robinson, C. T., & Gessner, M. O. (2000). Nutrient addition accelerates leaf breakdown in an alpine springbrook. Oecologia, 122(2), 258–263. DOI: 10.1007/pl00008854.

Rosemond, A. D., Mulholland, P. J., & Brawley, S. H. (2000). Seasonally shifting limitation of stream periphyton: Response of algal populations and assemblage biomass and productivity to variation in light, nutrients, and herbivores. Canadian Journal of Fisheries and Aquatic Sciences, 57(1), 66–75. DOI: 10.1139/f99-181.

Rosemond, A. D., Reice, S. R., Elwood, J. W., & Mulholland, P. J. (1992). The effects of stream acidity on benthic invertebrate communities in the south-eastern United States. Freshwater Biology, 27(2), 193–209. DOI: 10.1111/j.1365-2427.1992.tb00533.x.

Rosseland, B. O. & Kroglund, F. (2011) Lessons from acidification and pesticides. In: Atlantic salmon ecology (eds Ø. Aas, S. Einum, A. Klemetsen & J. Skurdal), pp. 387–407. Wiley-Blackwell, Oxford, England.

Rosset, J., Barlocher, F., & Oertli, J. J. 1982. Decomposition of conifer needles and deciduous leaves in two Black Forest and two Swiss Jura streams. Internationale Revue der gesamten Hydrobiologie 67, 695–711.

Rounick, J. S., & Gregory, S. V. (1981). Temporal changes in periphyton standing crop during an unusually dry winter in streams of the western Cascades, Oregon. Hydrobiologia, 83, 197–205. DOI: 10.1007/BF00008284.

Sagar, P. M., & Glova, G. J. (1992). Invertebrate drift in a large, braided New Zealand river. Freshwater Biology, 27(3), 405–416. DOI: 10.1111/j.1365-2427.1992.tb00550.x.

Scrine, J., Jochum, M., Ólafsson, J. S., & O’Gorman, E. J. (2017). Interactive effects of temperature and habitat complexity on freshwater communities. Ecology and Evolution, 7(22), 9333–9346. DOI: 10.1002/ece3.3412.

Scruton, D. A., Pennell, C. J., Robertson, M. J., Clarke, K. D., Eddy, W., & McKinley, R. S. (2005). Telemetry studies of the passage route and entrainment of downstream migrating wild Atlantic salmon (*Salmo salar*) smolts at two hydroelectric installations on the Exploits River, Newfoundland, Canada. In M. T. Spedicato, G. Lembo, & G. Marmulla (eds.), Aquatic telemetry: Advances and applications. Proceedings of the Fifth Conference on Fish Telemetry held in Europe, Ustica, Italy, 9–13 June 2003 (pp. 91–99). FAO/COISPA.

Shearer, K. A., Hayes, J. W., & Stark, J. D. (2002). Temporal and spatial quantification of aquatic invertebrate drift in the Maruia River, South Island, New Zealand. New Zealand Journal of Marine and Freshwater Research, 36(3), 529–536. DOI: 10.1080/00288330.2002.9517108.

Shortreed, K. S., & Stockner, J. G. (1983). Periphyton biomass and species composition in a coastal rainforest stream in British Columbia: Effects of environmental changes caused by logging. Canadian Journal of Fisheries and Aquatic Sciences, 40, 1887–1895. DOI: 10.1139/f83-219.

Simon, K. S., Chadwick, M. A., Huryn, A. D., & Valett, H. M. (2010). Stream ecosystem response to chronic deposition of N and acid at the Bear Brook Watershed, Maine. Environmental Monitoring and Assessment, 171(1), 83–92. DOI: 10.1007/s10661-010-1532-2.

Slobodkin, L. B. & Bossert, P. E. (2010) Cnidaria. In: Ecology and classification of North American freshwater invertebrates (eds J. H. Thorp & A. P. Covich), pp. 125–142. Academic Press, San Diego, CA. DOI: 10.1016/B978-0-12-374855-3.00005-4.

Sokol’skaya, N. L. (1975). A new species of Stylodrilus (Oligochaeta, Lumbriculidae) from the Chukchi Peninsula. Zoologicheskii Zhurnal, 54, 116–119.

Stals, R. (2015) Water beetles: Order Coleoptera. Chapter 14. In: Freshwater life: A field guide to the plants and animals of southern Africa (eds C. Griffiths, J. Day & M. Picker), pp. 194–209. Struik Nature, Cape Town.

Stovall, J. P., Keeton, W. S., & Kraft, C. E. (2009). Late-successional riparian forest structure results in heterogeneous periphyton distributions in low-order streams. Canadian Journal of Forest Research, 39(12), 2343–2354. DOI: 10.1139/X09-137.

Strommer, J. L., & Smock, L. A. (1989). Vertical distribution and abundance of invertebrates within the sandy substrate of a low-gradient headwater stream. Freshwater Biology, 22(2), 263–274. DOI: 10.1111/j.1365-2427.1989.tb01099.x.

Symons, P .E. K. (1979) Estimated escapement of Atlantic salmon (*Salmo salar*) for maximum smolt production in rivers of different productivity. Journal of the Fisheries Research Board of Canada, 36(2), 132–140. DOI: 10.1139/f79-022.

Tauber, C. A., Tauber, M. J., & Albuquerque, G. S. (2003) Neuroptera (Lacewings, Antlions). In: Encyclopedia of insects (eds V. H. Resh & R. T. Cardé), pp. 785–798. Academic Press, San Diego, CA.

Tonkin, J. D., Death, R. G., & Barquín, J. (2014). Periphyton control on stream invertebrate diversity: Is periphyton architecture more important than biomass? Marine and Freshwater Research, 65(9), 818–829. DOI: 10.1071/MF13271.

Vannote, R. L., Minshall, G. W., Cummins, K. W., Sedell, J. R., & Cushing, C. E. (1980). The river continuum concept. Canadian Journal of Fisheries and Aquatic Sciences, 37(1), 130–137. DOI: 10.1139/f80-017.

Veliz, M. A. (1999) Relationship of nutrients, and mayfly (Ephemeroptera) abundance and diet to periphyton biomass in boreal streams. Master’s Thesis, University of Alberta.

Vieira, N. K. M., Poff, N. L., Carlisle, D. M., Moulton, S. R., Koski, M. L., & Kondratieff, B. C. (2006). A database of lotic invertebrate traits for North America (Data Series 187). Reston, VA: U.S. Geological Survey.

Vinarski, M. V. (2019) Class Gastropoda. In: Thorp and Covich’s freshwater invertebrates: Keys to Palaearctic fauna (4th ed., Vol. 4) (eds D. C. Rogers & J. H. Thorp), pp. 310–345. Elsevier, London.

Von Schiller, D., Martí, E., Riera, J. L., & Sabater, F. (2007). Effects of nutrients and light on periphyton biomass and nitrogen uptake in Mediterranean streams with contrasting land uses. Freshwater Biology, 52(5), 891–906. DOI: 10.1111/j.1365-2427.2007.01742.x.

Voshell, J. R. (2002) A guide to common freshwater invertebrates of North America. McDonald & Woodward Publishing Company, Blacksburg, VA.

Wang, L., Robertson, D. M., & Garrison, P. J. (2007). Linkages between nutrients and assemblages of macroinvertebrates and fish in wadeable streams: Implication to nutrient criteria development. Environmental Management, 39(2), 194–212. DOI: 10.1007/s00267-006-0135-8.

Warnaars, T. A., Hondzo, M., & Power, M. E. (2007). Abiotic controls on periphyton accrual and metabolism in streams: Scaling by dimensionless numbers. Water Resources Research, 43(8), W08425. DOI: 10.1029/2006WR005002.

Warren, D. R., Keeton, W. S., Bechtold, H. A., & Rosi-Marshall, E. J. (2013). Comparing streambed light availability and canopy cover in streams with old-growth versus early-mature riparian forests in western Oregon. Aquatic Sciences, 75(4), 547–558. DOI: 10.1007/s00027-013-0299-2.

Wetzel, R. G. (ed.) (1983) Periphyton of freshwater ecosystems. Developments in Hydrobiology, 17. Dr W. Junk Publishers, The Hague. 346 pp.

Wootton, J. T. (2012). River food web response to large-scale riparian zone manipulations. PLoS One, 7(12), e51839. DOI: 10.1371/journal.pone.0051839.

