## Supporting Information for "Investigating biotic, chemical, and physical drivers of periphyton and invertebrate biomass in boreal streams supporting juvenile Atlantic salmon"

**2. Methods**

*2.2.1 Invertebrate biomass*

*2.2.1.a Invertebrate functional feeding groups*

In conjunction with invertebrate biomass calculations, numerous sources were used (Barbour, Gerritsen, Snyder, & Stribling, 1999; Tauber, Tauber, & Albuquerque, 2009; Oliveira & Nessimian, 2010; Dhooria, 2016; Collins III, 2017) to assign functional feeding groups (filter/collector, gatherer/collector, predator, scraper, shredder) to all identified invertebrate taxa to assess relative functional feeding group differences. See Table S7 for complete functional feeding group percentage contribution to overall invertebrate biomass.

*2.2.2 Periphyton biomass*

The following equation outlines the components used to calculate chlorophyll *a*. The before acidification step measures absorbance of both chlorophyll *a* and any additional pigments in the solution. The acidification step converts chlorophyll *a* to pheophyton *a* and provides an updated absorbance reading. The difference between the two absorbance readings in conjunction with other equation components is then used to estimate chlorophyll *a*.

$$Chlorophyll a (\mu g/{cm}^{2}) = \frac{26.7(E_{664b}- E_{665a}) \times Vext}{area of substrate ({cm}^{2}) \times L (cm)}$$

where *E_664b_* = [(absorbance of solution at 664 nm − absorbance of blank at 664 nm) − (absorbance of solution at 750 nm − absorbance of blank at 750 nm)] before acidification; *E_665b_* = [(absorbance of solution at 665 nm − absorbance of blank at 665 nm) − (absorbance of solution at 750 nm − absorbance of blank at 750 nm)] after acidification;

*V_ext_* = volume of 90% acetone used in the extraction (mL);

L = cuvette length (cm); and 26.7 = absorbance correction.

**Supporting information figures**

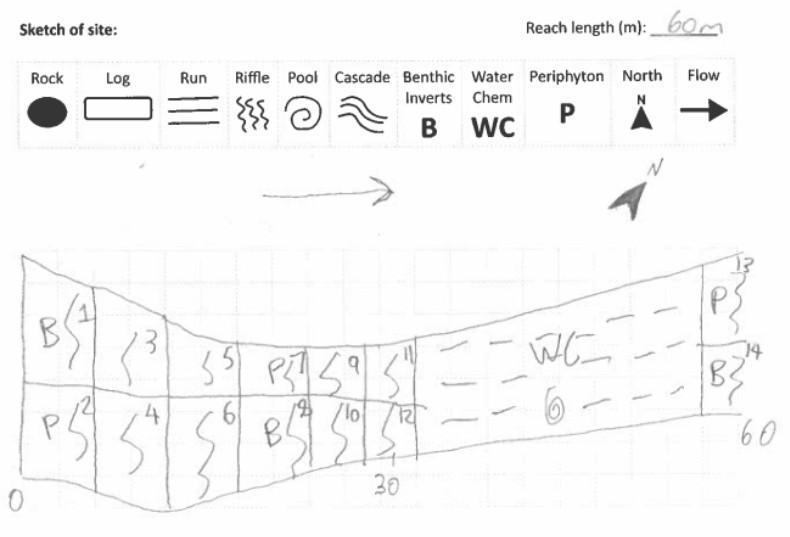

**Figure S1.** An example of how we mapped each site to determine Surber (B) and periphyton (P) collection locations. Water chemistry collection occurred in the farthest upstream run (WC). Site sketch from Charles Brook (CHB-CB-1) site (ID 257 in Figure 2).

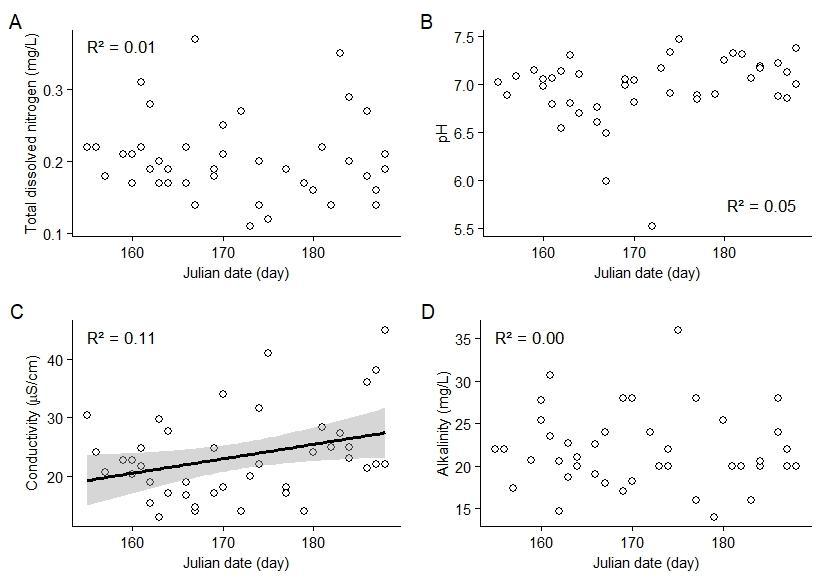

**Figure S2.** Relationships between (A) total dissolved nitrogen and Julian date, (B) pH and Julian date, (C) conductivity and Julian date, and (D) alkalinity and Julian date. Coefficient of determination (R²); linear regression lines and 95% confidence intervals (shaded) are shown only for statistically significant relationships (*p* < 0.05).

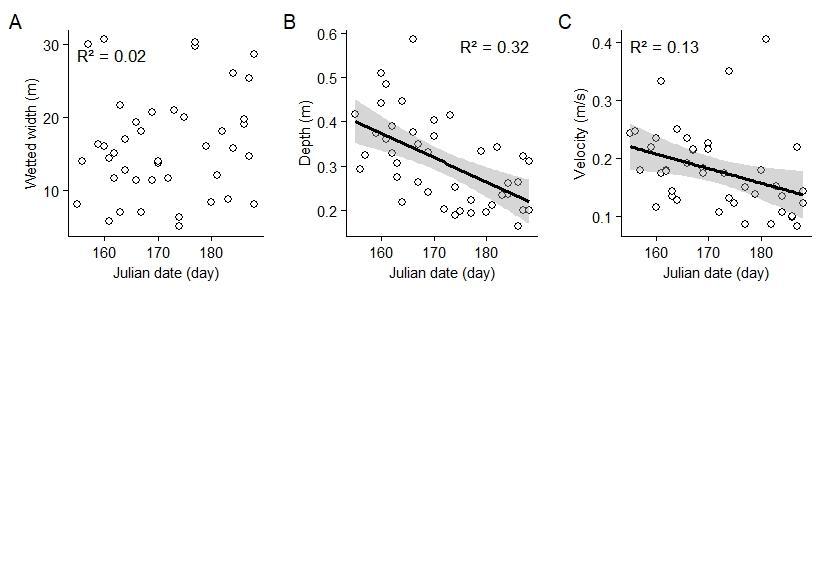

**Figure S3.** Relationships between (A) wetted width and Julian date, (B) depth and Julian date, and (C) velocity and Julian date. Coefficient of determination (R²); linear regression lines and 95% confidence intervals (shaded) are shown only for statistically significant relationships (*p* < 0.05).

**Supporting information tables**

**Table S1.** Date and location of sites and data logger deployment status. Site IDs correspond with IDs in Figure 2.

| Site ID | Site name | Date sampled | Latitude, longitude (decimal degrees) | Logger deployed (yes/no) |
| --- | --- | --- | --- | --- |
| 1 | Jumpers Brook | 4-Jun | 49.02148, -55.39841 | No |
| 5 | Tote Brook Trib 3 | 18-Jun | 48.81928, -55.48830 | Yes |
| 8 | Haynes Lake | 12-Jun | 48.77399, -55.49959 | Yes |
| 14 | Rocky Pond Brook | 9-Jun | 48.74296, -55.52735 | Yes |
| 16 | Beaver Brook | 20-Jun | 48.63250, -55.73509 | Yes |
| 39 | Paradise Lake Brook | 28-Jun | 48.70641, -55.63074 | Yes |
| 42 | Three Brook Trib | 7-Jun | 48.91119, -55.53818 | Yes |
| 45 | Little Rattling Brook | 18-Jun | 48.92155, -55.61128 | Yes |
| 49 | Green Woods | 3-Jun | 48.91731, -55.64837 | No |
| 51 | West Stoney Brook | 12-Jun | 48.85021, -55.78121 | Yes |
| 53 | Upper Stoney Brook | 30-Jun | 48.81482, -55.75401 | No |
| 67 | Lower Tom Joe Brook | 9-Jun | 48.92738, -55.95863 | Yes |
| 69 | Pamehoc Brook 1 | 14-Jun | 48.94124, -56.04375 | Yes |
| 72 | Mary Ann Brook 2 | 14-Jun | 49.07818, -55.98991 | Yes |
| 79 | Rocky Brook | 5-Jun | 49.06115, -56.11103 | No |
| 80 | Little Red Indian Trib 1 | 8-Jun | 48.92012, -56.19001 | Yes |
| 88 | Trappers Brook 3 | 10-Jun | 48.65672, -56.19354 | Yes |
| 106 | Snowshoe Pond Brook 1 | 15-Jun | 48.41224, -56.73853 | Yes |
| 111 | Michaels Brook | 15-Jun | 48.76747, -56.38759 | Yes |
| 112 | East Pond Brook 1 | 1-Jul | 48.68184, -56.51056 | No |
| 119 | South Steady | 10-Jun | 48.61371, -56.63494 | Yes |
| 121 | Harpoon Brook 5 | 2-Jul | 48.60433, -56.63440 | No |
| 125 | Burnt Pond Brook | 2-Jul | 48.74096, -56.59937 | No |
| 129 | Joe Glodes 3 | 8-Jun | 48.86519, -56.43868 | Yes |
| 143 | Little Sandy Brook 2 | 11-Jun | 48.80870, -56.69218 | Yes |
| 147 | Buchans Brook Trib 1B | 25-Jun | 48.84710, -56.78572 | No |
| 149 | Buchans Brook Trib 2B | 25-Jun | 48.85772, -56.82264 | No |
| 158 | Wileys Brook A | 17-Jun | 48.77128, -56.86586 | Yes |
| 163 | Skidder Pond Brook | 29-Jun | 48.73891, -56.93018 | No |
| 164 | Healeys Bay Brook | 11-Jun | 48.68560, -56.87787 | Yes |
| 170 | Half Way Brook 1 | 27-Jun | 48.46024, -56.79755 | No |
| 213 | Victoria River Trib 1A | 21-Jun | 48.42500, -57.08096 | No |
| 231 | Costigan Brook | 17-Jun | 48.55548, -57.11979 | Yes |
| 232 | Tulks Brook A | 23-Jun | 48.54099, -57.17885 | No |
| 253 | Lloyds River Tributary 1 | 22-Jun | 48.46067, -57.43143 | Yes |
| 255 | Lloyds River Tributary 2A | 22-Jun | 48.39334, -57.46984 | No |
| 257 | Charles Brook | 6-Jul | 49.34121, -55.25380 | No |
| 258 | Mill Pond Brook | 5-Jul | 49.40417, -55.47977 | No |
| 259 | Northern Arm Brook | 4-Jul | 49.08813, -55.59076 | No |
| 260 | New Bay River | 6-Jul | 49.24845, -55.40518 | No |
| 261 | Peters River | 4-Jul | 49.01959, -55.58184 | No |
| 262 | Western Arm Brook | 5-Jul | 49.34749, -55.46791 | No |

**Table S2.** Data collected across biotic, chemical, physical, temporal, and temperature categories, with variable names and units.

| Category | Variable | Units |
| --- | --- | --- |
| Biotic | Benthic invertebrates | g/m² |
|  | Periphyton | mg/cm² |
| Chemical | pH | pH units |
|  | Conductivity | μS/cm |
|  | Alkalinity | mg/L CaCO_3_ |
|  | Total dissolved nitrogen | mg/L |
|  | Phosphorus | ppm |
|  | Dissolved organic carbon | mg/L |
|  | Total dissolved solids | ppm |
| Physical | Wetted width | m |
|  | Depth | m |
|  | Velocity | m/s |
| Temporal | Julian date | Day |
| Temperature | Water temperature | °C |

**Table S3.** Candidate models used to investigate variation in invertebrate and periphyton biomass across sites. Candidate models consist of the following explanatory variables: periphyton biomass (Per_bio), invertebrate biomass (Invert_bio), total dissolved nitrogen (TDN), pH, conductivity (EC), alkalinity (Alk), wetted width (Wet_wid), depth (Depth), velocity (Velocity), and Julian date (Jul_d). Our modelling approach involved fitting models with biotic, chemical, physical, and temporal predictors and using model selection to select top ranking models per category.

| **Response** | **Category** | **Model number** | **Model** |
| --- | --- | --- | --- |
| **Invertebrate biomass** | **Biotic** | 1 | Null |
|  |  | 2 | Per_bio |
|  | **Chemical** | 1 | Null |
|  |  | 2 | TDN |
|  |  | 3 | pH |
|  |  | 4 | EC |
|  |  | 5 | Alk |
|  |  | 6 | TDN + pH |
|  |  | 7 | TDN + EC |
|  |  | 8 | TDN + Alk |
|  |  | 9 | TDN + pH + EC |
|  |  | 10 | TDN + pH + Alk |
|  |  | 11 | TDN + EC + Alk |
|  |  | 12 | pH + EC |
|  |  | 13 | pH + Alk |
|  |  | 14 | pH + EC + Alk |
|  |  | 15 | EC + Alk |
|  |  | 16 | TDN + Ph + EC + Alk |
|  | **Physical** | 1 | Null |
|  |  | 2 | Wet_wid |
|  |  | 3 | Depth |
|  |  | 4 | Velocity |
|  |  | 5 | Wet_wid + Depth |
|  |  | 6 | Wet_wid + Velocity |
|  |  | 7 | Depth + Velocity |
|  |  | 8 | Wet_wid + Depth + Velocity |
|  | **Temporal** | 1 | Null |
|  |  | 2 | Jul_d |
| **Periphyton biomass** | **Biotic** | 1 | Null |
|  |  | 2 | Invert_bio |
|  | **Chemical** | 1 | Null |
|  |  | 2 | TDN |
|  |  | 3 | pH |
|  |  | 4 | EC |
|  |  | 5 | Alk |
|  |  | 6 | TDN + pH |
|  |  | 7 | TDN + EC |
|  |  | 8 | TDN + Alk |
|  |  | 9 | TDN + pH + EC |
|  |  | 10 | TDN + pH + Alk |
|  |  | 11 | TDN + EC + Alk |
|  |  | 12 | pH + EC |
|  |  | 13 | pH + Alk |
|  |  | 14 | pH + EC + Alk |
|  |  | 15 | EC + Alk |
|  |  | 16 | TDN + Ph + EC + Alk |
|  | **Physical** | 1 | Null |
|  |  | 2 | Wet_Wid |
|  |  | 3 | Depth |
|  |  | 4 | Velocity |
|  |  | 5 | Wet_Wid + Depth |
|  |  | 6 | Wet_Wid + Velocity |
|  |  | 7 | Depth + Velocity |
|  |  | 8 | Wet_Wid + Depth + Velocity |
|  | **Temporal** | 1 | Null |
|  |  | 2 | Jul_d |

**Table S4.** Candidate models used to investigate variation in invertebrate and periphyton biomass across sites. Candidate models consist of the following explanatory variables: average spring water temperature (Spring_temp), average summer water temperature (Summer_temp), average fall water temperature (Fall_temp), and average winter water temperature (Winter_temp). Our modelling approach involved fitting models with temperature predictors and using model selection to select top ranking models.

| **Response** | **Category** | **Model number** | **Model** |
| --- | --- | --- | --- |
| **Invertebrate biomass** | **Temperature** | 1 | Null |
|  |  | 2 | Spring_temp |
|  |  | 3 | Summer_temp |
|  |  | 4 | Fall_temp |
|  |  | 5 | Winter_temp |
| **Periphyton biomass** | **Temperature** | 1 | Null |
|  |  | 2 | Spring_temp |
|  |  | 3 | Summer_temp |
|  |  | 4 | Fall_temp |
|  |  | 5 | Winter_temp |

**Table S5.** Results of generalized linear models using Gamma error distributions and log link functions to explain invertebrate and periphyton biomass (*n* = 42). We present all models in each category (i.e., biotic, chemical, physical, and temporal). *K*: the number of model parameters, AIC_c_: Akaike’s Information Criterion corrected for small sample sizes model value, ∆AIC_c_: change in Akaike’s Information Criterion corrected for small sample sizes relative to top model, AIC_c_ weight: Akaike’s Information Criterion corrected for small sample sizes model weight, LL: log likelihood of model, R^2^: Nagelkerke’s pseudo R^2^ (the proportion of variation in data explained by the model). Models that are italicized have uninformative variables (see Leroux, 2019).

| **Model parameter** | ***K*** | **AIC_c_** | **ΔAIC_c_** | **AIC_c_ weight** | **LL** | **R^2^** |
| --- | --- | --- | --- | --- | --- | --- |
| Invertebrate biomass |  |  |  |  |  |  |
| Biotic |  |  |  |  |  |  |
| Periphyton biomass | 3 | 188.98 | 0.00 | 0.80 | -91.18 | 0.15 |
| Intercept | 2 | 191.81 | 2.83 | 0.20 | -93.75 | 0.00 |
| Chemical |  |  |  |  |  |  |
| Conductivity | 3 | 191.19 | 0.00 | 0.20 | -92.28 | 0.07 |
| Conductivity + Alkalinity | 4 | 191.79 | 0.61 | 0.15 | -91.36 | 0.11 |
| Intercept | 2 | 191.81 | 0.62 | 0.15 | -93.75 | 0.00 |
| Alkalinity | 3 | 193.25 | 2.06 | 0.07 | -93.31 | 0.03 |
| *pH + Conductivity* | 4 | 193.46 | 2.27 | 0.06 | -92.19 | 0.07 |
| *TDN + Conductivity* | 4 | 193.59 | 2.40 | 0.06 | -92.25 | 0.07 |
| *pH* | 3 | 193.62 | 2.43 | 0.06 | -93.49 | 0.01 |
| *pH + Conductivity + Alkalinity* | 5 | 194.05 | 2.87 | 0.05 | -91.19 | 0.11 |
| *TDN* | 3 | 194.13 | 2.95 | 0.05 | -93.75 | 0.00 |
| *TDN + Conductivity + Alkalinity* | 5 | 194.30 | 3.12 | 0.04 | -91.32 | 0.11 |
| *pH + Alkalinity* | 4 | 195.10 | 3.91 | 0.03 | -93.01 | 0.04 |
| *TDN + Alkalinity* | 4 | 195.69 | 4.51 | 0.02 | -93.31 | 0.03 |
| *TDN + pH* | 4 | 195.99 | 4.81 | 0.02 | -93.46 | 0.02 |
| *TDN + pH + Conductivity* | 5 | 196.04 | 4.86 | 0.02 | -92.19 | 0.07 |
| *TDN + pH + Conductivity + Alkalinity* | 6 | 196.78 | 5.60 | 0.01 | -91.19 | 0.11 |
| *TDN + pH + Alkalinity* | 5 | 197.61 | 6.42 | 0.01 | -92.97 | 0.04 |
| Physical |  |  |  |  |  |  |
| Wetted width + Velocity | 4 | 191.17 | 0.00 | 0.22 | -91.04 | 0.13 |
| Velocity | 3 | 191.71 | 0.54 | 0.17 | -92.54 | 0.06 |
| Intercept | 2 | 191.81 | 0.64 | 0.16 | -93.75 | 0.00 |
| Wetted width | 3 | 191.90 | 0.74 | 0.15 | -92.64 | 0.07 |
| *Depth* | 3 | 192.81 | 1.65 | 0.10 | -93.09 | 0.03 |
| *Wetted width + Depth* | 4 | 193.30 | 2.13 | 0.08 | -92.11 | 0.08 |
| *Wetted width + Depth + Velocity* | 5 | 193.60 | 2.43 | 0.07 | -90.96 | 0.12 |
| *Depth + Velocity* | 4 | 193.70 | 2.54 | 0.06 | -92.31 | 0.06 |
| Temporal |  |  |  |  |  |  |
| Julian date | 3 | 186.46 | 0.00 | 0.94 | -89.91 | 0.20 |
| Intercept | 2 | 191.81 | 5.35 | 0.06 | -93.75 | 0.00 |
| Periphyton biomass |  |  |  |  |  |  |
| Biotic |  |  |  |  |  |  |
| Invertebrate biomass | 3 | -167.73 | 0.00 | 1 | 87.18 | 0.33 |
| Intercept | 2 | -156.08 | 11.65 | 0 | 80.19 | 0.00 |
| Chemical |  |  |  |  |  |  |
| Alkalinity | 3 | -162.21 | 0.00 | 0.26 | 84.42 | 0.18 |
| TDN + Alkalinity | 4 | -162.02 | 0.18 | 0.23 | 85.55 | 0.17 |
| *pH + Alkalinity* | 4 | -160.84 | 1.37 | 0.13 | 84.96 | 0.18 |
| *Conductivity + Alkalinity* | 4 | -159.83 | 2.38 | 0.08 | 84.46 | 0.17 |
| *TDN + pH + Alkalinity* | 5 | -159.58 | 2.63 | 0.07 | 85.62 | 0.17 |
| *TDN + Conductivity + Alkalinity* | 5 | -159.45 | 2.75 | 0.06 | 85.56 | 0.17 |
| TDN | 3 | -158.46 | 3.74 | 0.04 | 82.55 | 0.11 |
| *pH + Conductivity + Alkalinity* | 5 | -158.46 | 3.75 | 0.04 | 85.06 | 0.19 |
| *TDN + pH + Conductivity + Alkalinity* | 6 | -157.03 | 5.18 | 0.02 | 85.72 | 0.18 |
| *TDN + Conductivity* | 4 | -157.01 | 5.20 | 0.02 | 83.04 | 0.14 |
| Intercept | 2 | -156.08 | 6.13 | 0.01 | 80.19 | 0.00 |
| *TDN + pH* | 4 | 156.01 | 6.19 | 0.01 | 82.55 | 0.11 |
| *TDN + pH + Conductivity* | 5 | -154.98 | 7.23 | 0.01 | 83.32 | 0.14 |
| *pH + Conductivity* | 4 | -154.77 | 7.43 | 0.01 | 81.93 | 0.06 |
| *pH* | 3 | -154.61 | 7.60 | 0.01 | 80.62 | 0.02 |
| *Conductivity* | 3 | -154.44 | 7.77 | 0.01 | 80.54 | 0.01 |
| Physical |  |  |  |  |  |  |
| Velocity | 3 | -158.96 | 0.00 | 0.45 | 82..80 | 0.10 |
| *Wetted width + Velocity* | 4 | -156.69 | 2.27 | 0.15 | 82.88 | 0.10 |
| *Depth + Velocity* | 4 | -156.58 | 2.38 | 0.14 | 82.83 | 0.10 |
| Intercept | 2 | -156.08 | 2.88 | 0.11 | 80.19 | 0.00 |
| *Wetted width* | 3 | -154.58 | 4.39 | 0.05 | 80.60 | 0.02 |
| *Depth* | 3 | -154.30 | 4.66 | 0.04 | 80.47 | 0.01 |
| *Wetted width + Depth + Velocity* | 5 | -154.17 | 4.79 | 0.04 | 82.92 | 0.10 |
| *Wetted width + Depth* | 4 | -152.58 | 6.38 | 0.02 | 80.83 | 0.03 |
| Temporal |  |  |  |  |  |  |
| Julian date | 3 | -157.95 | 0.00 | 0.72 | 82.29 | 0.08 |
| Intercept | 2 | -156.08 | 1.87 | 0.28 | 80.19 | 0.00 |

**Table S6.** Results of generalized linear models using Gamma error distributions and log link functions to explain invertebrate and periphyton biomass (*n* = 22). We present all models in the temperature category. *K*: the number of model parameters, AIC_c_: Akaike’s Information Criterion corrected for small sample sizes model value, ∆AIC_c_: change in Akaike’s Information Criterion corrected for small sample sizes relative to top model, AIC_c_ weight: Akaike’s Information Criterion corrected for small sample sizes model weight, LL: log likelihood of model, R^2^: Nagelkerke’s pseudo R^2^ (the proportion of variation in data explained by the model).

| **Model parameter** | ***K*** | **AIC_c_** | **ΔAIC_c_** | **AIC_c_ weight** | **LL** | **R^2^** |
| --- | --- | --- | --- | --- | --- | --- |
| Invertebrate biomass |  |  |  |  |  |  |
| Fall water temperature | 3 | 93.22 | 0.00 | 0.43 | -42.94 | 0.18 |
| Intercept | 2 | 94.33 | 1.12 | 0.25 | -44.85 | 0.00 |
| Winter water temperature | 3 | 95.63 | 2.42 | 0.13 | -44.15 | 0.07 |
| Spring water temperature | 3 | 95.76 | 2.54 | 0.12 | -44.21 | 0.07 |
| Summer water temperature | 3 | 97.03 | 3.82 | 0.06 | -44.85 | 0.00 |
| Periphyton biomass |  |  |  |  |  |  |
| Summer water temperature | 3 | -102.06 | 0.00 | 0.34 | 54.70 | 0.18 |
| Fall water temperature | 3 | -101.41 | 0.64 | 0.25 | 54.37 | 0.10 |
| Intercept | 2 | -101.08 | 0.98 | 0.21 | 52.86 | 0.00 |
| Winter water temperature | 3 | -99.74 | 2.32 | 0.11 | 53.54 | 0.04 |
| Spring water temperature | 3 | -99.47 | 2.59 | 0.09 | 53.40 | 0.04 |

**Table S7.** Complete functional feeding group percentage contribution to overall invertebrate biomass from invertebrates collected and taxonomically identified. Note, percentages do not sum to 100%, as functional feeding groups could not be assigned to immature Plecoptera and Trichoptera.

| Functional feeding group | Gatherer/collector | Filterer/collector | Predator | Scraper (grazer) | Shredder |
| --- | --- | --- | --- | --- | --- |
| Percentage (%) contribution to overall invertebrate biomass | 56.58 | 26.48 | 7.62 | 6.13 | 2.93 |

**Supporting information box**
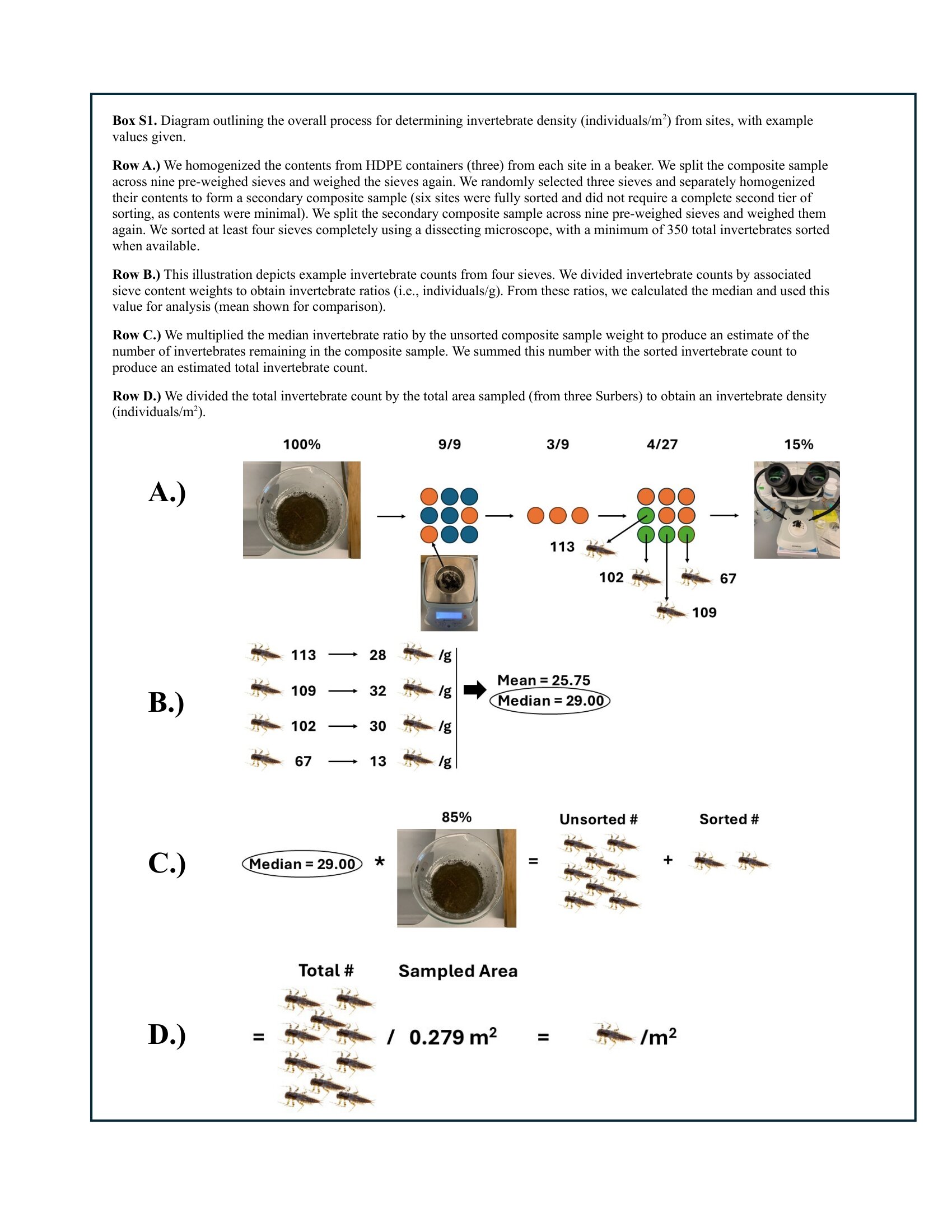
